# Contrasting patterns and co-occurrence network of soil bacterial and fungal community along depth profiles in cold temperate montane forests of China

**DOI:** 10.1101/2021.04.11.439312

**Authors:** Li Ji, Fangyuan Shen, Yue Liu, Yuchun Yang, Jun Wang, Lixue Yang

**Affiliations:** Key Laboratory of Sustainable Forest Ecosystem Management-Ministry of Education, School of Forestry, Northeast Forestry University, Harbin 150040, P.R. China; Jilin Academy of Forestry, Changchun 130033, P.R. China

**Keywords:** soil profile, bacteria, fungi, cold temperate forest, co-occurrence network, Illumina MiSeq sequencing

## Abstract

Soil bacterial and fungal communities with different key ecological functions play an important role in the boreal forest ecosystem. Despite several studies have reported the microbial altitudinal distribution patterns, our understanding about the characteristics of the microbial community and the core composition of the microbiome in cold-temperate mountain forests is still limited. In this study, Illumina MiSeq sequencing was used to investigate the changes in soil bacterial and fungal communities in surface and subsurface soils along at an altitudinal gradient (from 830 m to 1300 m) on Oakley Mountain in the northern Greater Khingan Mountains. Altitude and soil depth had significant impacts on the relative abundance of Proteobacteria, Acidobacteria and Actinobacteria (dominant phylum for bacteria), and altitude had significant impacts on the Ascomycota, Basidiomycota and Mucoromycota (dominant phylum for fungi). The diversity of bacterial and fungal communities showed a monotonous decrease and increase with altitude. The influence of altitude on bacterial and fungal community composition was greater than that of soil depth. The variation of pH and dissolved organic nitrogen (DON) content in different altitudes were the main factors driving the bacterial and fungal community structure, respectively. There is no obvious difference between the network structure of surface and subsurface soil fungal communities, while the network of subsurface soil bacterial communities was more complex and compact than the surface layer. The network nodes mainly belonging to Proteobacteria and Actinobacteria are the key species in the two soil layers. Our results demonstrated that the altitude had a stronger influence on soil bacterial and fungal communities than soil depth, and bacterial and fungal communities showed divergent patterns along the altitudes and soil profiles.

## 1 Introduction

Soil microorganisms are an important part of the forest ecosystem, and play a critical role in nutrient conversion, organic matter decomposition and energy flow. The altitude distribution pattern of soil microorganisms is one of the important contents of the biogeographic distribution pattern and has been ignored for a long time. In recent years, with the development of sequencing technology, many scholars focus on the biogeography of soil microbes, and found that the soil microbial community is monotonously decreasing (Bryant et al., 2008; Bahram et al., 2012; Shen et al., 2019), humpback (Miyamoto et al., 2014; Li et al., 2016; Peay et al., 2017) or on significant patterns along the altitudinal gradients (Fierer et al., 2011; Shen et al., 2014). However, these studies are mostly concentrated in tropical, subtropical and temperate regions, and only limited studies have been reported on the altitudinal distribution of soil microbial communities in cold temperate regions (Jarvis et al., 2015). The cold temperate forests are considered to be an important habitat for storing a large amount of biomass carbon (about 30%), have low productivity and nutrient cycling rate (Reich, 2012), which is very sensitive to climate change, especially in soil microorganisms and biochemical cycle processes (Christensen et al., 2004; Reich et al., 2012). The rapid response of microorganisms to changes in environmental conditions and their high turnover rate may provide more information that indicates ecosystem service (Banning et al., 2011).

Soil bacteria and fungi, as important components of the microflora, exert important ecological functions. Given that their different morphological characteristics, growth rate, environmental sensitivity, phylogeny and life history, they exhibit divergent biogeographic patterns (Hannula et al., 2017). Some studies reported that the growth rate of soil bacteria was approximately ten times higher than that of specific soil fungi, and soil fungi tend to be more resistant to low-temperature soil habitats (Rousk and Bååth, 2007; Kirchman, 2018). In a recent study, Ma et al. (2017) found soil bacteria and fungi had a unique biogeographic distribution in forest soils across continental-scale, and dispersal limitation and environmental variables dominated the variation of bacterial and fungal communities. In the mountain ecosystem, the high variability of plant communities and soil properties along the altitudinal gradient inevitably leads to dramatic variations for bacteria and fungi. Jarvis et al. (2015) found that temperature was the main factor affecting the ectomycorrhizal fungal community in the Mt. Cairngorm in Scotland. The results of Singh et al. (2014) in Mt. Halla also proved that the annual average temperature and precipitation played a critical role in the changes in the structure and composition of bacterial communities along the altitudinal gradient. Moreover, some scholars believed that temperature had a positive correlation with the species richness of animals, plants and microorganisms (Hawkins et al., 2003; Zhou et al., 2016). The metabolic theory based on ecology explained this temperature-diversity relationship. The biochemical kinetics of metabolism predicted that biodiversity increases with increasing temperature (Brown et al., 2004). Compared to tropical and subtropical regions, there is still uncertainty in the cold temperate regions with specific climatic conditions, however, whether soil bacterial and fungal communities have obvious altitude distribution patterns, and whether temperature or other environmental factors dominate this variation.

In addition to abiotic factors, biotic factors (interactions among species) are also considered to be the complementary mechanism that affects the biogeographic patterns of microorganisms (Fan et al., 2017). The symbiosis, parasitism, competition or predation among different microorganisms in the community will form a complex co-occurrence network (Faust and Raes, 2012). In recent years, utilizing network analysis, numerous studies had reported on the interaction and biological complexity of soil microorganisms in forest ecosystems (Xiao et al., 2018; Li et al., 2020; Tu et al., 2020). The topological characteristics of the network are used to evaluate keystone species that regulated ecosystem function and community stability (Cardona et al., 2016). Some recent studies had indicated the relationship between vertical distribution and interaction of microbial communities along the soil profiles from the perspective of network analysis (Yang et al., 2017; Luan et al., 2020). To date, most studies only focused on the changes in the microbial community and related processes in surface soil regarding altitudinal gradient (Eilers et al., 2012; Sheng et al., 2019), however, far too little attention had been paid to subsurface soil, microorganisms in subsurface soil play a key role in soil formation and biogeochemical cycle processes, and exhibit greater variation and characteristics different from surface soil (Fritze et al., 2000). Soil depth can increase the rate of microbial evolution, including gene mutation, community assembly and interaction, and the microbial community has high stability in the upper soil, while the opposite occurs in the subsurface soil (Du et al., 2021). Some studies had pronounced the variation in soil physicochemical properties affected the microbial diversity and community composition along different soil depths in harsh climate areas (Coolen et al., 2011; Deng et al., 2015). However, due to the complexity of the soil microbiome, much less is known about the interactions between microbial members in the community, which limits our understanding of their role in ecosystem functions (Widder et al., 2016). As far as we know, the information on the keystones in the soil microbial communities in the boreal forest ecosystem is still limited. The composition of these microbial communities along the soil profiles and the framework affecting their community assembly are yet to be explored.

Mountain ecosystems, as an important component of terrestrial ecosystems, which the regulating services of forests are of particular importance (Seidl et al., 2019). The habitats have a wide variety of habitats with rapidly changing climate, vegetation and soil quality can be found in harsh mountain environments (Sundqvist et al., 2013), these regulating for forests services are of particular importance. Oakley mountain is the highest peak in the northern in Greater Khingan Mountain, relative to 1520 m above sea level, however, the soil microbial distribution pattern of the boreal forest ecosystem dominated by larch is rarely reported, which limits our prediction of the response of soil microbial community to climate change in the cold temperate region. Given that the high variability of soil microbiomes along an altitudinal gradient and the fundamental differences in life strategies between bacteria and fungi (Baldrian, 2017), here, we compared the diversity and co-occurrence networks of soil bacterial and fungal communities along an altitudinal gradient in cold temperate, and it will generate fresh insight into the main ecological predictors of microbiology along altitudinal gradients. We hypothesized that (1) with increasing altitude, the diversity, and structure of soil bacterial and fungal community will show consistent patterns, i.e. monotonical decline; (2) given previous findings on the factors that dominate changes in the bacterial and fungal community, temperature and pH may be key factors affecting changes in the composition and structure of fungi and bacterial community in cold temperate, respectively; and (3) the soil bacterial and fungal communities inhabiting surface and subsurface soil will perform different network topology characteristics.

## 2 Materials and methods

### 2.1 Study area and sampling

The study area was on Oakley Mountain (51°50’N, 122°01’E) at the Pioneer Forest Farm of the A’longshan Forestry Bureau in the northern Greater Khingan Mountains in China. This area is an ideal location for investigating the biogeographical patterns of soil microbes along an altitudinal gradient due to the steep topography. This area is characterized by a cold temperate climate with long, cold winters and short, warm summers. The annual mean air temperature is −5.1°C, and the annual mean precipitation is 437.4 mm. Oakley Mountain is the highest mountain in the northern Greater Khingan Mountains and is covered by snow from October to May, i.e., for approximately 7 months each year (Figs. S1). The soils are mostly Umbri-Gelic Cambosols according to the Chinese taxonomic system, with an average depth of 20 cm.

Based on the variation in the composition of the vegetation community along the altitudinal gradient (Table S1), soil samples were collected from the southern slope of Oakley Mountain at four sites representing different elevations (830, 950, 1100, and 1300 m). At each site, we created three 20 × 30 m plots, and eight soil cores of 0~10 cm and 10~20 cm soil depth were randomly collected and thoroughly mixed to make a composite sample for each plot. The soil samples were collected in July (mid-growing season) 2019 (N=24) and immediately transported on ice to the laboratory. We used a button temperature sensor (HOBO H8 Pro, Onset Complete Corp., Bourne, MA, USA) to record the soil temperature (ST) of each plot. The fresh soil samples were sieved through a 2 mm sieve, and visible roots and other residues were removed. Each sample was divided into two subsamples: one that was stored at −80 °C for DNA extraction, and one that was stored at 4 °C for the measurement of soil physicochemical properties. Basic information about the sites at the different elevations is provided in Supplementary Table S1.

### 2.2 Soil physicochemical properties

The soil moisture (SM) and bulk density (BD) were measured by the cutting ring method. The soil pH was measured using a pH meter (MT-5000, Shanghai) after shaking a soil water (1:5 w/v) suspension for 30 min. The soil organic carbon (SOC) and total nitrogen (TN) contents in each sample were measured after tableting using a J200 Tandem laser spectroscopic element analyzer (Applied Spectra, Inc., Fremont, CA, USA), and the total phosphorus (P) content was determined colorimetrically with a UV spectrophotometer (TU-1901, Puxi Ltd., Beijing, China) after wet digestion with HClO_4_-H_2_SO_4_. The soil dissolved organic carbon (DOC) content was analyzed using a total organic carbon (TOC) analyzer (Analytik Jena, Multi N/C 3000, Germany), and the soil nitrate (NO_3_^-^-N), ammonium (NH_4_^+^-N), and total dissolved nitrogen (DTN) contents were determined using a continuous flow analytical system (AA3, Seal Co., Germany). The soil dissolved organic nitrogen (DON) was calculated from the soil contents of NO_3_^-^-N, NH_4_^+^-N, and DTN. Soil microbial biomass carbon and nitrogen were determined by the chloroform fumigation method (Brookes et al., 1985; Joergensen, 1996).

### 2.3 DNA extraction and PCR amplification

Soil bacterial and fungal DNA was extracted from the soil samples using an E.Z.N.A.^®^ Soil DNA Kit (Omega Biotek, Norcross, GA, U.S.) based on the manufacturer’s protocols. The concentration and purity of the DNA were measured with a NanoDrop 2000 UV-vis spectrophotometer (Thermo Scientific, Wilmington, USA). The quality and quantity of the extracted DNA were evaluated with a 1.0% (w/v) agarose gel. The bacterial 16S and fungal ITS genes were amplified. For bacteria, the primers 338F (5’-ACTCCTACGGGAGGCAGCAG-3’) and 806R (5’-GGACTACHVGGGTWTCTAAT-3’) were used to amplify the V3-V4 region. For fungi, the primers ITS3F (5’-GCATCGATGAAGAACGCAGC-3’) and ITS4R (5’-TCCTCCGCTTATTGATATGC-3’) were used to amplify the ITS2 region (Lee et al., 2012; Gade et al., 2013). All bacterial and fungal primers were performed with a thermocycler PCR system (GeneAmp 9700, ABI, USA). PCRs were performed in triplicate in a 20 μL mixture composed of 4 μL of 5× FastPfu Buffer, 2 μL of 2.5 mM dNTPs, 0.8 μL of each primer (5 μM), 0.4 μL of FastPfu Polymerase and 10 ng of template DNA. The PCRs were conducted using the following program: 3 min of denaturation at 95 °C; 30 cycles of 30 s at 95 °C, 30 s for annealing at 55 °C, and 45 s for elongation at 72 °C; and a final extension at 72 °C for 10 min (Caporaso et al., 2012).

### 2.4 Illumina MiSeq sequencing and processing of the sequencing data

A 2% agarose gel was used to extract the PCR products, which were then purified with the AxyPrep DNA Gel Extraction Kit (Axygen Biosciences, Union City, CA, USA). Based on the manufacturer’s protocols, the products were quantified using a QuantiFluor™-ST fluorometer (Promega, USA). Subsequently, the amplicons were merged on the Illumina MiSeq platform (Illumina, San Diego, USA) in equimolar amounts and paired-end sequenced (2 × 300 bp) following the standard protocols of Majorbio Bio-Pharm Technology Co., Ltd. (Shanghai, China). We deposited the raw reads into the NCBI database (accession number: PRJNA721110 for bacteria, PRJNA721105 for fungi).

Trimmomatic was used to demultiplex the raw fastq files and conduct quality filtering, and the reads were merged with FLASH with the following procedures implemented (Caporaso et al., 2012): (i) The read segments with an average quality score <20 in a 50 bp sliding window were truncated. (ii) Primers were precisely matched, allowing mismatches between two nucleotides, and read segments with ambiguous bases were deleted. (iii) Sequences with an overlap length of more than 10 bp were combined according to their overlapping portion. UPARSE (version 7.1, http://drive5.com/uparse/) was used to cluster the sequences into operational taxonomic units (OTUs) based on 97% similarity (Edgar, 2013), and UCHIME was used to identify chimeric sequences. The taxonomic identities of the gene sequences for each 16S and ITS were assigned by BLAST against the SILVA bacterial and the UNITE fungal ITS database, respectively.

### 2.5 Co-occurrence network analysis

Based on random matrix theory (RMT), the molecular ecological network analysis method (http://ieg4.rccc.ou.edu/mena/) was used to construct networks for the different soil profiles. In most cases, only nodes detected in half or more of the total sample were retained for subsequent network construction. For more information about related theories, algorithms and procedures, please refer to Deng et al. (2012) and Zhou et al. (2011). Spearman rank correlation was used to establish a co-occurrence network among the soil bacterial and fungal communities, respectively. When constructing the network, the same similarity threshold (*St*) was used to ensure that the co-occurrence networks in different seasons could be compared with each other. Then, the same network size and average number of links were used to generate 100 corresponding random networks. The *Z*-test was used to test for differences between the empirical network and the random networks. The intra-module connectivity value (*Zi*) and inter-module connectivity value (*Pi*) for each node were used to identify the keystone species in the network (Deng et al., 2016; Olesen et al., 2007). In this study, we used the following simplified classification and evaluation criteria: (i) peripheral nodes (*Zi*≤2.5, *Pi*≤0.62), which have only a few links that almost always connect to nodes in their modules; (ii) highly linked connector nodes (*Zi*≤2.5, *Pi*> 0.62), which have many modules; (iii) module nodes (*Zi*> 2.5, *Pi*≤0.62), which are highly connected to many nodes in their respective modules; and (iv) network nodes (*Zi*>2.5, *Pi*>0.62), which act as both module nodes and connection nodes. To show the results more clearly, Cytoscape (version 3.7.1) was used to visualize the co-occurrence networks of the soil bacteria and fungi (Cline et al., 2007).

### 2.6 Data analysis

The alpha diversity indices (observed number of OTUs (Sobs), Chao1, Faith’s phylogenetic diversity (PD), and Simpson) of the Illumina MiSeq sequencing data were analyzed with QIIME (Caporaso et al., 2012). The Shapiro-Wilk test and Levene test were used to evaluate the normality of the data and the homogeneity of variance. Nonmetric multidimensional scaling analysis (NMDS) of beta diversity based on Bray-Curtis distances was conducted with the ‘vegan’ package in R (version 3.6.1) to analyze bacterial and fungal community similarity. Analysis of similarities (ANOSIM) and permutation multivariate analysis of variance (PERMANOVA) of the Bray-Curtis distances were conducted to test for differences in the properties of the soil bacterial and fungal communities among different altitudes and depths. To identify the effects of soil properties on the soil bacterial and fungal communities, redundancy analysis (RDA) was used to predict the variation in the communities, and a Mantel test with a Monte Carlo simulation consisting of 999 randomizations was performed. The RDA function of the ‘vegan’ package and the mantel.rtest function of the ‘ade4’ package in R were used to perform the RDA and Mantel test, respectively (version 3.6.1) (Team, 2013).

## 3 Results

### 3.1 Soil physicochemical properties along altitudinal gradients

Altitude had significant effects on soil bulk density (BD), soil moisture (SM), temperature (ST), pH, inorganic nitrogen, MBC, MBN and DON (P<0.05, Table 1). As the altitude increases, the BD showed a significant decreasing trend, and the SM showed a significant increasing trend. ST and soil pH at 830m was significantly lower and higher than other altitudes, respectively. Soil nitrate nitrogen and ammonium nitrogen are the highest at 1300m and 1100m, respectively. MBC was the highest at 1100 m, which was 242.78% higher than 830m, and MBN was the highest at 950 m, which was 274.64% higher than 830m. Soil depth had a significant effect on SM and ST (*P*<0.05), and the SM and ST of the surface soil were significantly higher than those of the subsurface soil. The interaction of altitude and soil depth had no significant effect on all soil factors.

**Table 1.** The soil physicochemical property for surface and subsurface soils in different altitudes

| Soil variables | 830 m |  | 950 m |  | 1100 m |  | 1300 m |  | Two-way ANOVA |  |  |
| --- | --- | --- | --- | --- | --- | --- | --- | --- | --- | --- | --- |
|  | Surface soil | Subsurface soil | Surface soil | Subsurface soil | Surface soil | Subsurface soil | Surface soil | Subsurface soil | Altitude | Depth | A×D |
| BD (g·cm<sup>-3</sup>) | 0.89±0.13Aa | 1.18±0.16Aa | 0.63±0.10ABa | 0.81±0.08Ba | 0.58±0.03ABa | 0.64±0.04Ba | 0.34±0.02Ba | 0.45±0.04Ba | *** | ns | ns |
| SM (%) | 30.75±3.48Ca | 20.34±1.31Cb | 37.48±1.75BCa | 30.7±3.41BCb | 47.14±3.85Ba | 41.71±5.89Bb | 71.01±3.96Aa | 58.32±3.55Ab | *** | ** | ns |
| ST (□) | 9.01±0.87Ba | 7.00±0.58Bb | 10.41±0.19ABa | 9.02±0.29Ab | 11.61±0.55Aa | 9.48±0.84Ab | 10.25±0.35ABa | 9.23±0.28Ab | ** | ** | ns |
| SOC (mg·g<sup>-1</sup>) | 61.28±2.65Aa | 62.08±3.75Aa | 65.62±3.26Aa | 62.03±0.23Ba | 63.14±1.45Aa | 61.22±1.91Ba | 61.67±2.54Aa | 63.83±2.62Aa | ns | ns | ns |
| TN (mg·g<sup>-1</sup>) | 7.02±0.18Aa | 6.87±0.32Aa | 6.64±0.30Aa | 6.97±0.01Aa | 6.88±0.15Aa | 7.06±0.12Aa | 6.97±0.17Aa | 6.75±0.20Aa | ns | ns | ns |
| TP (mg·g<sup>-1</sup>) | 0.63±0.03Aa | 0.56±0.03Aa | 0.47±0.04Aa | 0.62±0.04Aa | 0.44±0.04Aa | 0.34±0.07Aa | 0.52±0.07Ab | 1.59±0.17Aa | ns | ns | ns |
| pH | 4.68±0.19Aa | 4.74±0.36Aa | 3.92±0.10Ba | 4.21±0.18Aa | 4.23±0.04Ba | 4.28±0.06Aa | 4.01±0.63Ba | 4.23±0.07Aa | ** | ns | ns |
| NO<sub>3</sub><sup>-</sup>-N (mg·kg<sup>-1</sup>) | 5.39±0.08Ba | 5.35±0.25Ba | 5.92±0.18Ba | 5.62±0.04Ba | 5.57±0.09Ba | 5.39±0.24Ba | 6.97±0.32Aa | 6.43±0.31Aa | *** | ns | ns |
| NH<sub>4</sub><sup>+</sup>-N (mg·kg<sup>-1</sup>) | 63.25±5.23Ba | 63.34±5.25Ba | 61.27±2.32Ba | 58.59±0.53Ba | 94.68±4.35Aa | 87.72±2.44Aa | 88.09±4.84ABa | 82.02±3.88Aa | *** | ns | ns |
| DOC (mg·kg<sup>-1</sup>) | 163.33±9.58Ba | 148.13±4.80Ba | 269.60±17.07Aa | 295.87±9.12Aa | 291.47±15.44Aa | 262.80±11.31Aa | 213.07±9.18Aa | 233.33±12.41ABa | ns | ns | ns |
| DON (mg·kg<sup>-1</sup>) | 15.92±2.31Ba | 11.48±1.75Ba | 17.57±1.61Ba | 15.34±1.22Ba | 33.33±2.59Aa | 36.57±2.12Aa | 32.61±4.74Aa | 33.73±1.85Aa | *** | ns | ns |
| MBC (mg·kg<sup>-1</sup>) | 456.84±26.86Ba | 307.02±11.88Bb | 602.81±59.38Ba | 538.95±45.01Ba | 1565.96±98.72Aa | 1515.79±24.21Aa | 785.26±54.04Ba | 885.26±75.11ABa | * | ns | ns |
| MBN (mg·kg<sup>-1</sup>) | 16.05±1.36Ba | 10.11±1.79Ba | 60.13±5.40Aa | 34.69±1.89Ab | 37.78±1.86Ba | 36.23±1.15Aa | 34.29±1.15Ba | 34.59±1.53Aa | * | ns | ns |

A: Altitude; D: Soil depth. Data with different uppercase letters were significantly difference at 5% level among different altitudes in the same soil layer ( $P<0.05$ ), while different lowercase letters indicate significant differences among different soil layers in the same altitude ( $P<0.05$ ); ns, not significant; \*, $P<0.05$ ; \*\*, $P<0.01$ .

### 3.2 Soil bacterial and fungal sequencing summary and community composition

The 16S rRNA genes for soil bacteria and ITS genes for fungi were sequenced on the Illumina MiSeq platform. Across all soil samples analyzed, 1,472,023 high-quality soil bacterial and 1,527,911 high-quality soil fungal sequences were obtained by Illumina MiSeq sequencing, respectively. A total of 49674~74237 (mean = 61334) soil bacterial and 46434~74407 (mean = 63662) soil fungal sequences were obtained per sample. The average read length for bacteria and fungi were 411 bp and 317 bp, which were larger than 99% of Good’s coverage for the 16S and ITS gene regions, respectively. The rarefaction curves of the genes tended to approach the saturation plateau at 97% sequence similarity for all samples (Fig. S2), which indicated that the sequencing depth was adequate for evaluating the structure and diversity of soil bacteria and fungi across all samples.

For soil bacteria, a total of 6577 OTUs were identified, distributed in 31 phyla, 91 classes and 646 genera. Proteobacteria, Acidobacteria and Actinobacteria were the dominant phyla, accounting for 75.8% of the total number of bacterial sequences obtained (Fig. 1A). Altitude and soil depth had a significant effect on the relative abundance of Chloroflexi, Planctomycetes and Firmicutes (Table S2). Alphaproteobacteria, Acidobacteriia and Actinobacteria were the dominant classes, with the relative abundance of 27.0%, 20.0% and 18.2%, respectively (Figure 1B). The interaction between altitude and soil depth had no significant effect on the relative abundances of all bacterial phyla and classes (Table S2).

**Figure 1.**
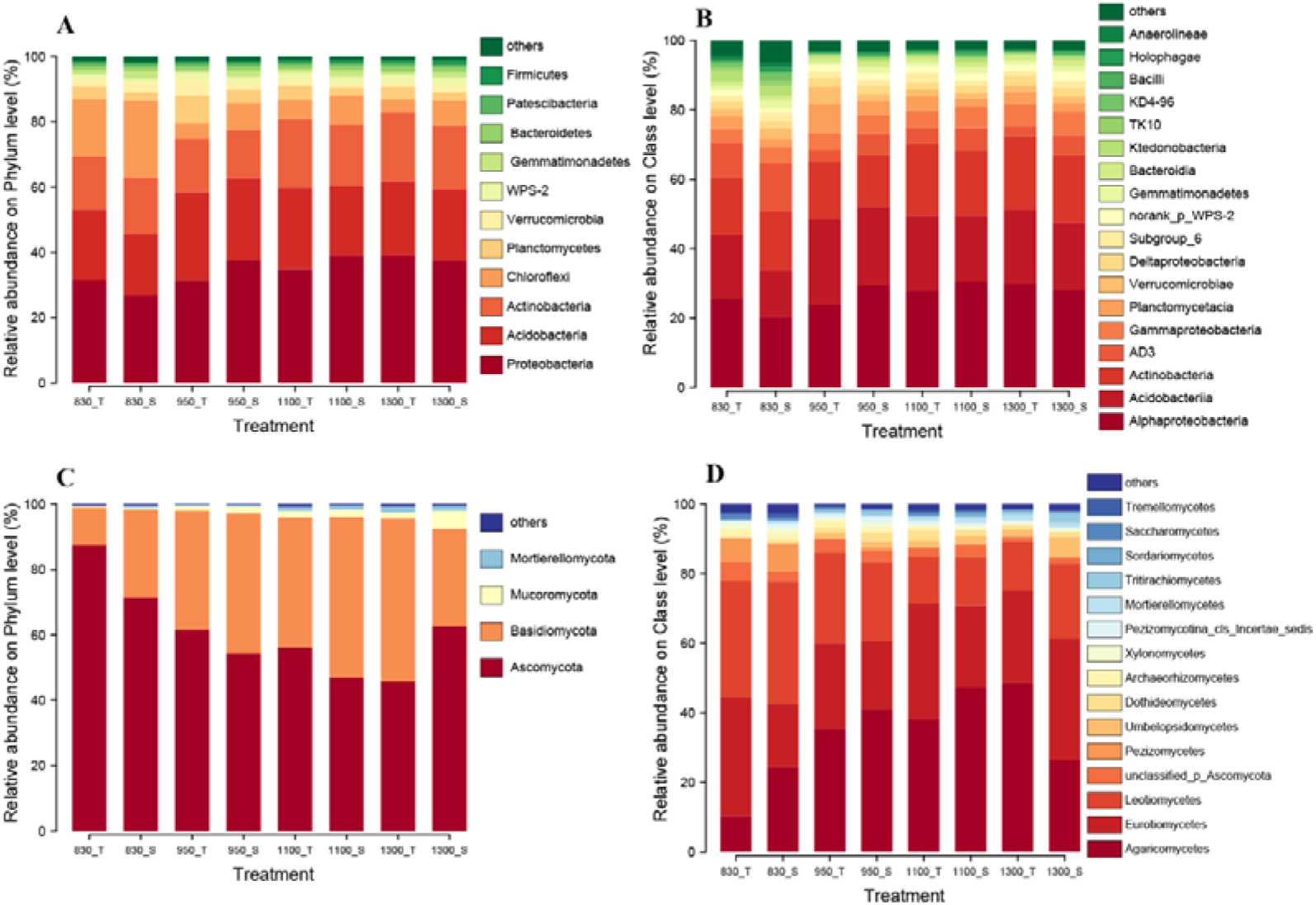
Relative abundances of main soil bacterial and fungal phyla (A, C) and classes (B, D) for surface and subsurface soils in different altitudes. 830_T, 950_T, 1100_T and 1300_T indicate the surface soil in 830 m, 950 m, 1100 m and 1300 m, respectively. 830_S, 950_S, 1100_S and 1300_S indicate the subsurface soil in 830 m, 950 m, 1100 m and 1300 m, respectively.

For soil fungi, a total of 2739 OTUs were identified, distributed in 14 phyla, 51 classes and 548 genera. At phylum level, fungal communities were dominated by Ascomycota and Basidiomycota, with the relative abundance of 60.8% and 35.8%, respectively (Figure 1C). Altitude had a marked effect on the Ascomycota, Basidiomycota and Mucoromycota, with increasing altitude, the relative abundance of Ascomycota showed a gradually decreasing trend (Table S3, Figure 1C). The dominant fungi were Agaricomycetes, Eurotiomycetes and Leotiomycetes at class level, and their relative abundance accounted for 83.4% of the total number of fungal sequences (Figure 1D). Soil depth, altitude and their interaction had no significant difference on the relative abundance of all fungal phyla and classes (Table S3).

### 3.3 Soil bacterial and fungal community diversity

Altitude had a significant impact on the Sobs, Chao1 and Faith’s PD diversity indices of soil bacterial communities (Figure 2A–2D). In general, the diversity of bacterial communities decreased with increasing altitude. In the 0-10 cm soil layer, the Sobs, Chao1 and Faith’s PD indices of soil bacteria at 830 m were 23.5%, 25.4% and 28.9% higher than those at 1300 m, respectively (*P*<0.05). In the 10-20 cm soil layer, the Sobs, Chao1 and Faith’s PD indices of soil bacterial at 830 m were 21.1%, 23.3% and 26.2% higher than those at 1300 m (*P*<0.05). The soil depth had no significant effect on the alpha diversity of the soil bacterial community.

**Figure 2.**
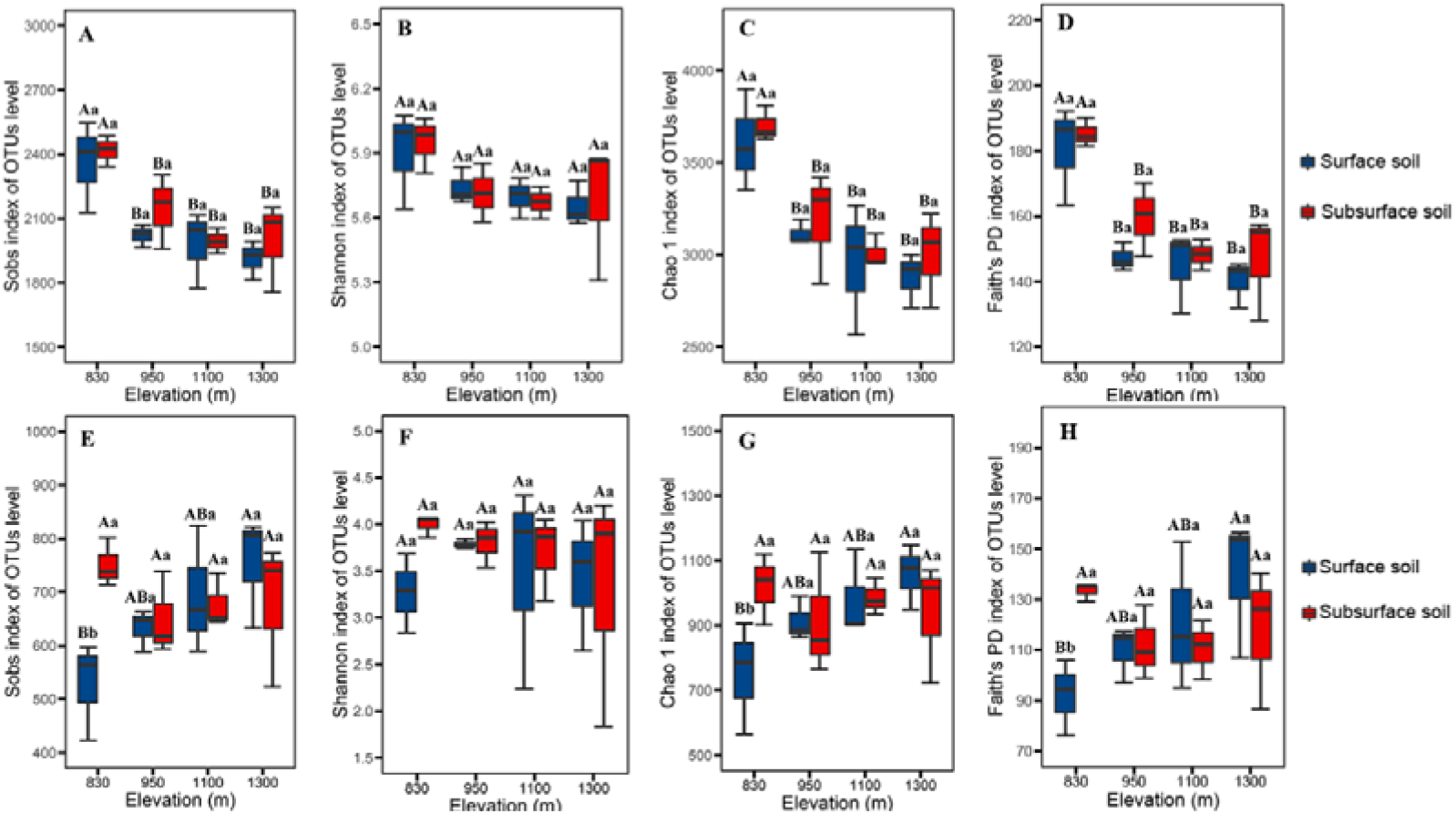
Sobs, Shannon, Chao1 and Faith’s PD indices of soil bacterial and fungal communities for surface and subsurface soils in different altitudes. OTUs were delineated at 97% sequence similarity. These indices were calculated using bacterial and fungal random subsamples of 49674 and 46343 gene sequences per sample. Two-way ANOVA for altitude and soil depth was conducted.

The fungal community alpha diversity index showed a potential increasing trend with altitude. In the 0-10 cm soil layer, the fungal Sobs, Chao1 and Faith’s PD indices at 1300 m were 42.7%, 40.6% and 50.8% higher than those at 830 m, respectively, while there was no significant difference in the 10-20 cm soil layer (Figure 2E, 2G and 2H). The soil depth had no significant effect on the alpha diversity of the fungal community (Figure 2E–2H).

NMDS analysis based on Bray-Curtis distance was performed on the soil bacterial and fungal sequencing data corresponding to the different altitudes for two contrasting soil depths. The bacterial and fungal community were divided into an obvious group based on altitude, while the group of soil depth in the same altitude for bacteria and fungi was not evident (Figure 3A and 3B). Compare to soil fungi, the bacterial community at 950 m, 1100 m and 1300 m were clustered and more similar. ANOSIM and PERMANOVA revealed significant differences in the structure of both soil bacterial and fungal community among altitudes (*P*<0.01, Figure 3). The PERMANOVA results of all samples demonstrated that altitude had a stronger influence than soil depth on the structure of the soil bacterial and fungal community (*P*<0.01, Table S4).

**Figure 3.**
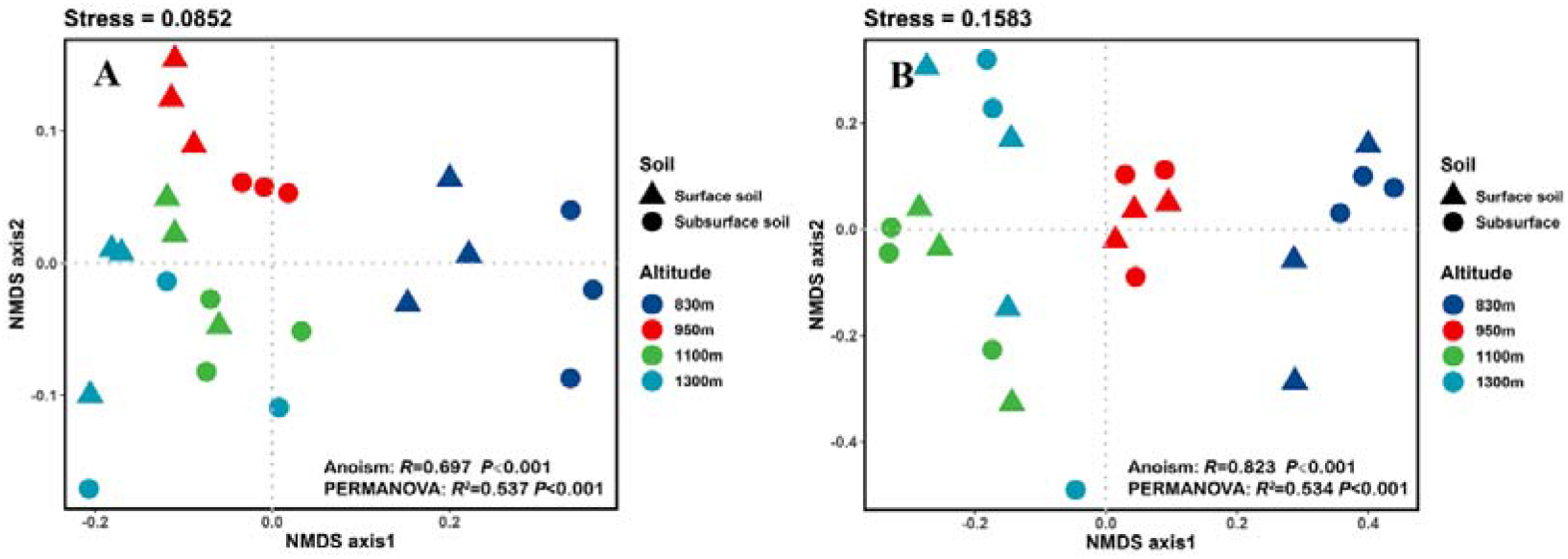
Principal coordinate analysis of soil bacterial (A) and fungal (B) communities based on Bray-Curtis distances.

### 3.4 Relationship between the soil microbial community and soil factors

The relationships between the soil factors and microbial community structure were evaluated by RDA and the Mantel test. The biplots showed that the first two axes explained more than 75.5% of the variation in both bacterial and fungal community structure (Figure 4A and 4B). However, there were differences in the main factors affecting bacterial and fungal community structure. For soil bacteria, pH was the main influencing factor, followed by BD and SM. Notably, DON exerted a significant effect on soil fungal community structure (Table 2).

**Figure 4.**
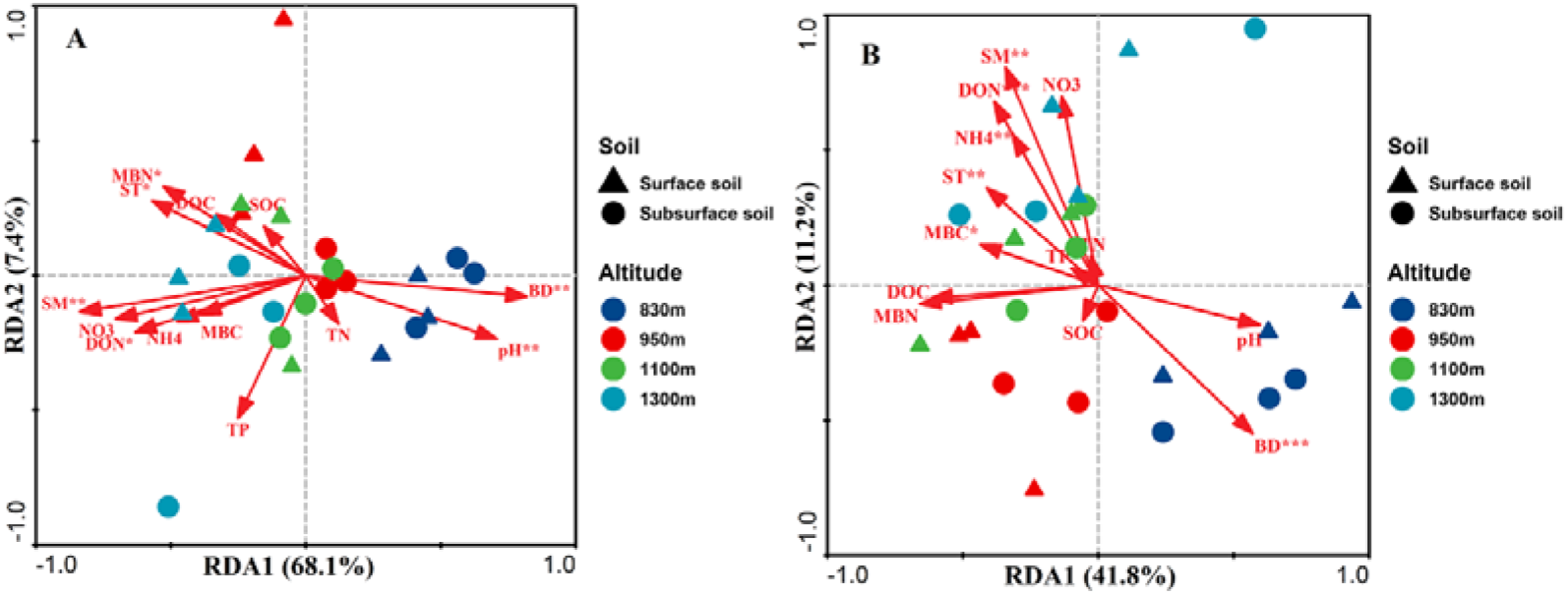
Redundancy analysis based on soil bacterial (A) and fungal (B) community at the genus level and soil factors (red arrows). The top 20 most abundant classified bacterial and fungal genera (97% sequence similarity) in the soil samples. Direction of arrow indicates the soil factors associated with changes in the community structure, and the length of the arrow indicates the magnitude of the association. The asterisk represents the significant soil factors associated with the bacterial or fungal community. The percentage of variation explained by RDA 1 and 2 is shown.

**Table 2.** Mantel test results for the correlation between relative abundance of bacterial and fungal genera and soil variables in different soil depths along the altitudinal gradient.

| Soil variables | Bacteria |  | Fungi |  |
| --- | --- | --- | --- | --- |
| | $R^2$ | $P$ | $R^2$ | $P$ |
| BD | 0.418 | **0.007** | 0.402 | **0.008** |
| SM | 0.379 | **0.008** | 0.551 | **0.002** |
| ST | 0.327 | **0.013** | 0.336 | **0.017** |
| SOC | 0.008 | 0.932 | 0.107 | 0.309 |
| TN | 0.005 | 0.959 | 0.132 | 0.224 |
| TP | 0.282 | 0.051 | 0.053 | 0.494 |
| pH | 0.419 | **0.002** | 0.232 | 0.077 |
| NO<sub>3</sub><sup>-</sup>-N | 0.229 | 0.070 | 0.237 | 0.068 |
| NH<sub>4</sub><sup>+</sup>-N | 0.197 | 0.093 | 0.501 | **0.003** |
| DOC | 0.179 | 0.118 | 0.054 | 0.571 |
| DON | 0.179 | **0.024** | 0.707 | **0.001** |
| MBC | 0.199 | 0.096 | 0.354 | **0.017** |
| MBN | 0.349 | **0.012** | 0.066 | 0.493 |

### 3.5 Soil bacterial and fungal co-occurrence patterns

The bacterial and fungal co-occurrence networks were constructed with different soil depths. For soil bacteria, the nodes of OTUs in the network mainly belonged to Proteobacteria, Actinobacteria, Chloroflexi and Acidobacteria, and the nodes of bacterial community are divided into 11 and 21 modules in the 0-10 cm and 10-20 cm soil layers, respectively (Figure 5A, 5B). Compared with the 0-10 cm soil layer, the number of nodes and connections of the bacterial community in the 10-20 cm soil layer increased significantly, and its network topological characteristics had a higher average degree, average clustering coefficient and average path length (Table 3). In the 0-10 cm and 10-20 cm soil layers, there had 92.9% and 90.8% of the proportions of positive interaction connections were observed, respectively. For fungi, most of the nodes belonged to Ascomycota and Basidiomycota, and generated 12 modules for each soil layer (Figure 6A, 6B). The two soil layers had similar number of nodes, links, average degree and average clustering coefficient, moreover, the positive and negative connections of the two soil layers were similar. Compared with the 0-10 cm soil layer, the 10-20 cm layer soil fungal network had a higher average path length and degree of modularity (Table 3).

**Figure 5.**
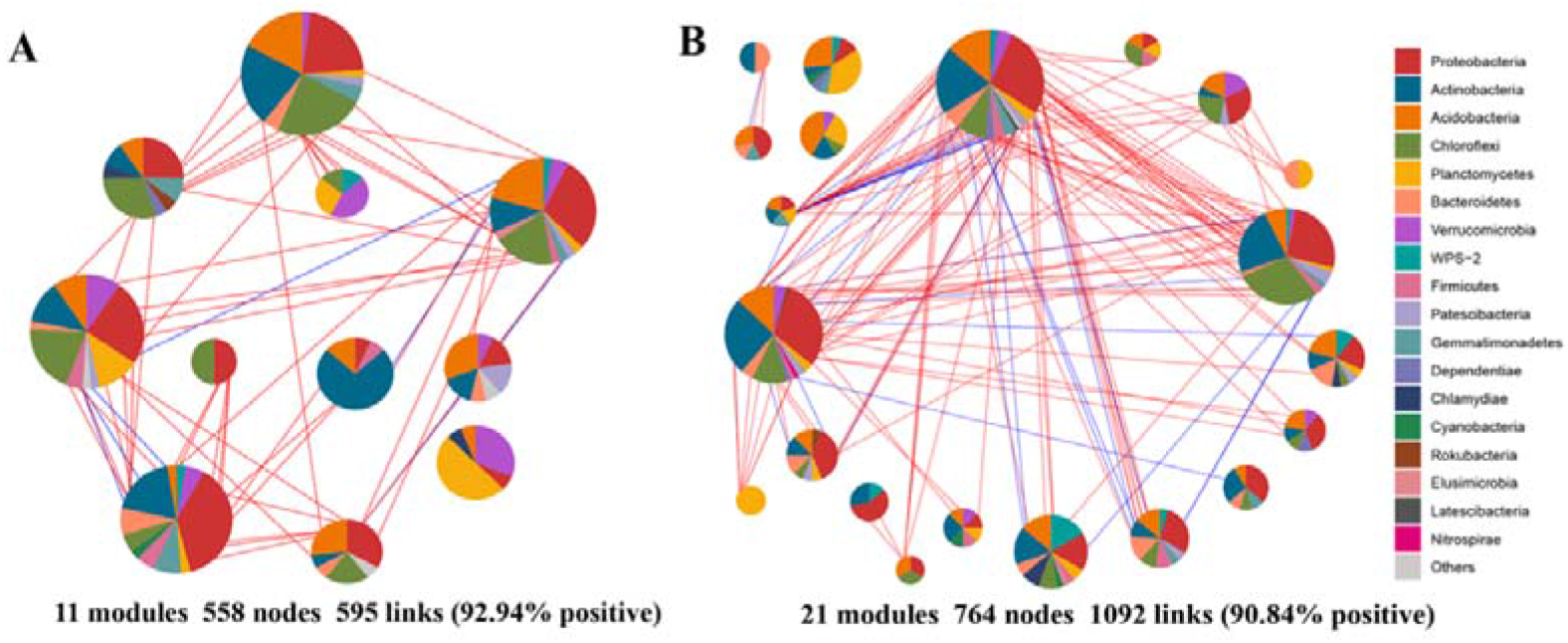
Overview of the co-occurrence networks for bacterial communities in surface and subsurface soils and the bacterial phylum-level composition of the dominant modules. Node size is proportional to the relative abundance. Major phylum (with nodes > 5) were randomly colored. Positive links between nodes were colored red and negative links were colored blue.

**Figure 6.**
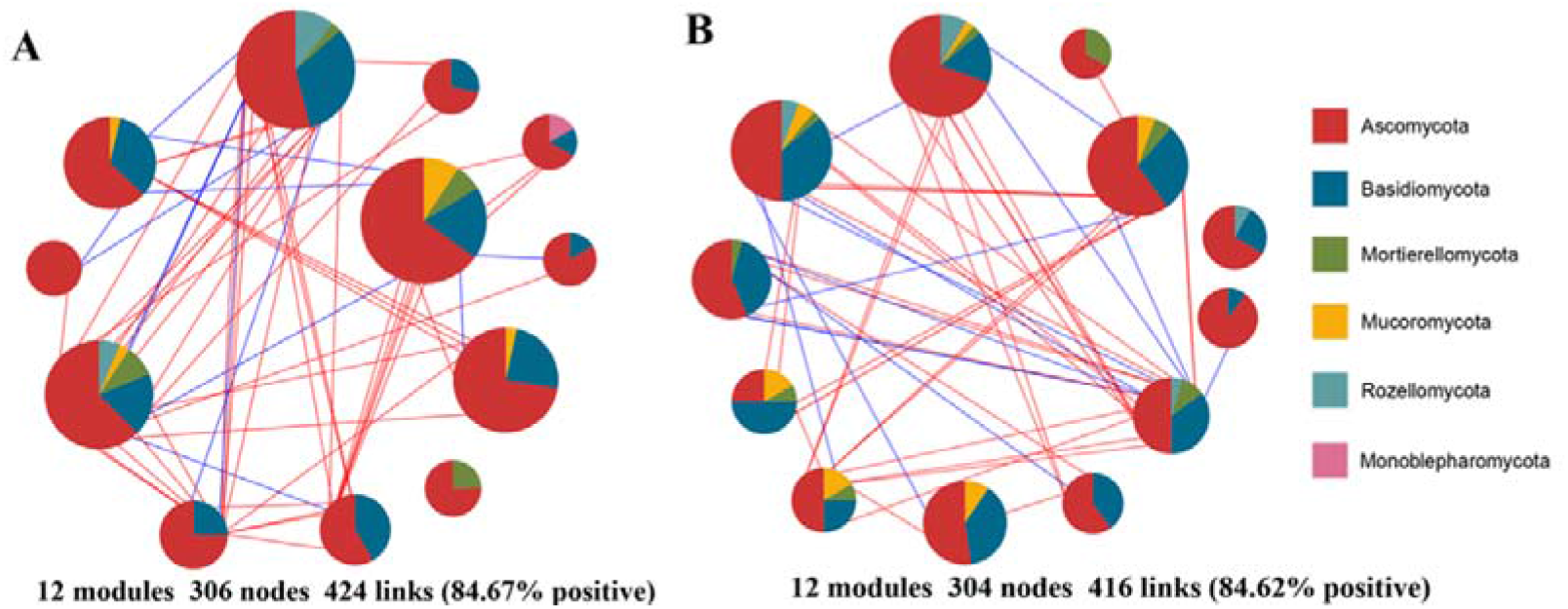
Overview of the co-occurrence networks for fungal communities in surface and subsurface soils and the fungal phylum-level composition of the dominant modules. Node size is proportional to the relative abundance. Major phylum (with nodes > 5) were randomly colored. Positive links between nodes were colored red and negative links were colored blue.

**Table 3.**
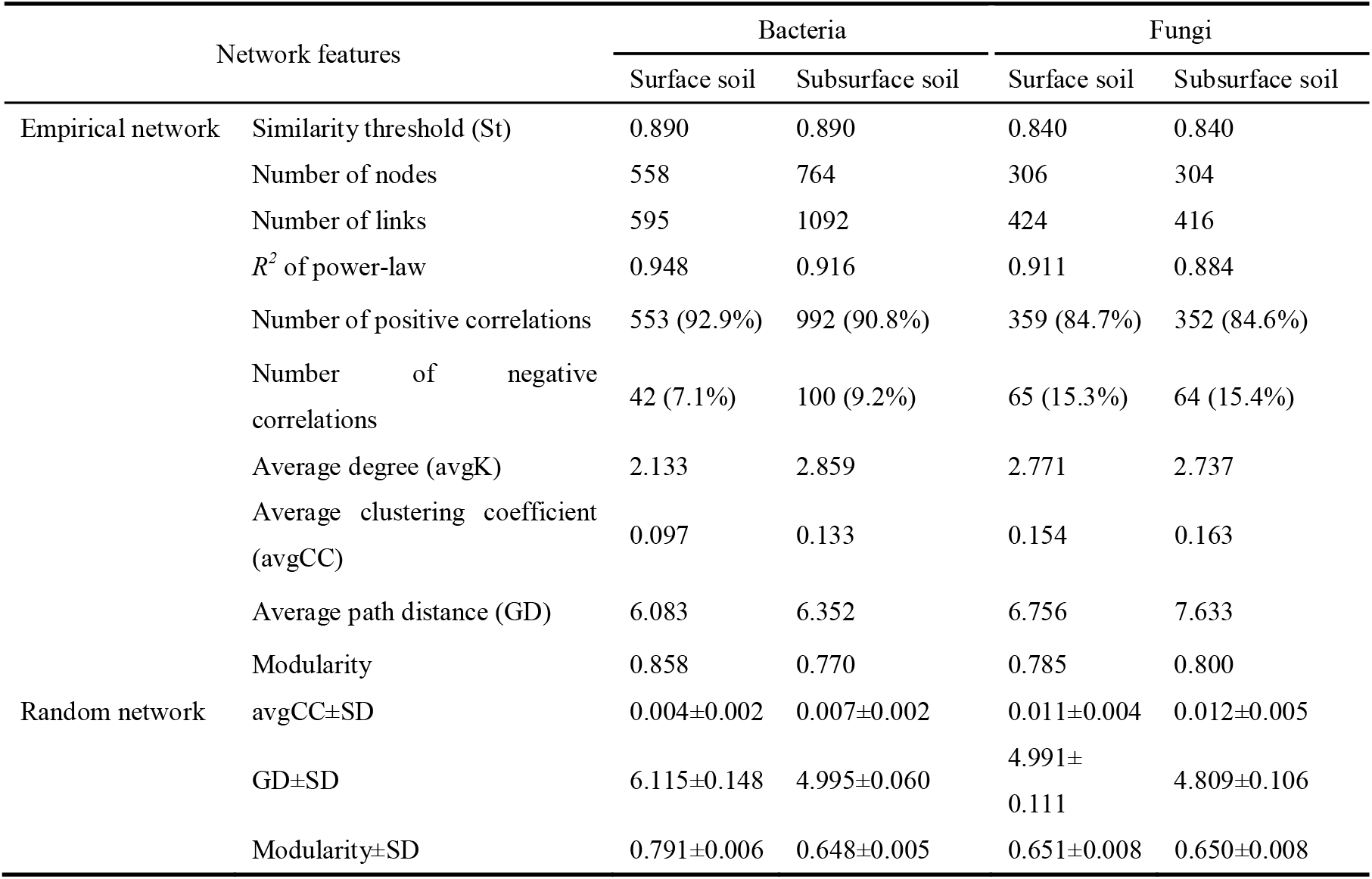
The topological properties for soil bacterial and fungal co-occurrence networks in different soil depths

| Network features |  | Bacteria |  | Fungi |  |
| --- | --- | --- | --- | --- | --- |
|  |  | Surface soil | Subsurface soil | Surface soil | Subsurface soil |
| Empirical network | Similarity threshold (St) | 0.890 | 0.890 | 0.840 | 0.840 |
|  | Number of nodes | 558 | 764 | 306 | 304 |
|  | Number of links | 595 | 1092 | 424 | 416 |
| | $R^2$ of power-law | 0.948 | 0.916 | 0.911 | 0.884 |
|  | Number of positive correlations | 553 (92.9%) | 992 (90.8%) | 359 (84.7%) | 352 (84.6%) |
|  | Number of negative correlations | 42 (7.1%) | 100 (9.2%) | 65 (15.3%) | 64 (15.4%) |
|  | Average degree (avgK) | 2.133 | 2.859 | 2.771 | 2.737 |
|  | Average clustering coefficient (avgCC) | 0.097 | 0.133 | 0.154 | 0.163 |
|  | Average path distance (GD) | 6.083 | 6.352 | 6.756 | 7.633 |
|  | Modularity | 0.858 | 0.770 | 0.785 | 0.800 |
| Random network | avgCC±SD | 0.004±0.002 | 0.007±0.002 | 0.011±0.004 | 0.012±0.005 |
| | GD±SD | 6.115±0.148 | 4.995±0.060 | 4.991±0.111 | 4.809±0.106 |
| | Modularity±SD | 0.791±0.006 | 0.648±0.005 | 0.651±0.008 | 0.650±0.008 |

Based on Zi and Pi values, we defined the peripheral, network connectors, module hubs and network hubs in the network. Zi-Pi scatter plots for all bacterial and fungal nodes in two contrasting soil layers were generated based on the module network. No node belonged to both the module hubs and the network connectors. There are 98.3% and 97.7% of the nodes as peripheral nodes in the bacterial and fungal networks, respectively, and they were highly connected in their respective modules (Figure 7A, 7B). For the bacterial network, 11 nodes (mainly belonged to Proteobacteria and Acidobacteria) were classified as module hubs, and they had strong associations with many nodes in their modules. There had 12 nodes were specifically classified as connectors between modules (Figure 7A). There had 10 nodes (belonging to Ascomycota and Basidiomycota) and 4 nodes (belonging to Ascomycota, Basidiomycota, and Mucoromycota) in the fungal network, respectively, which are classified as module hubs and network connectors (Figure 7B).

**Figure 7.**
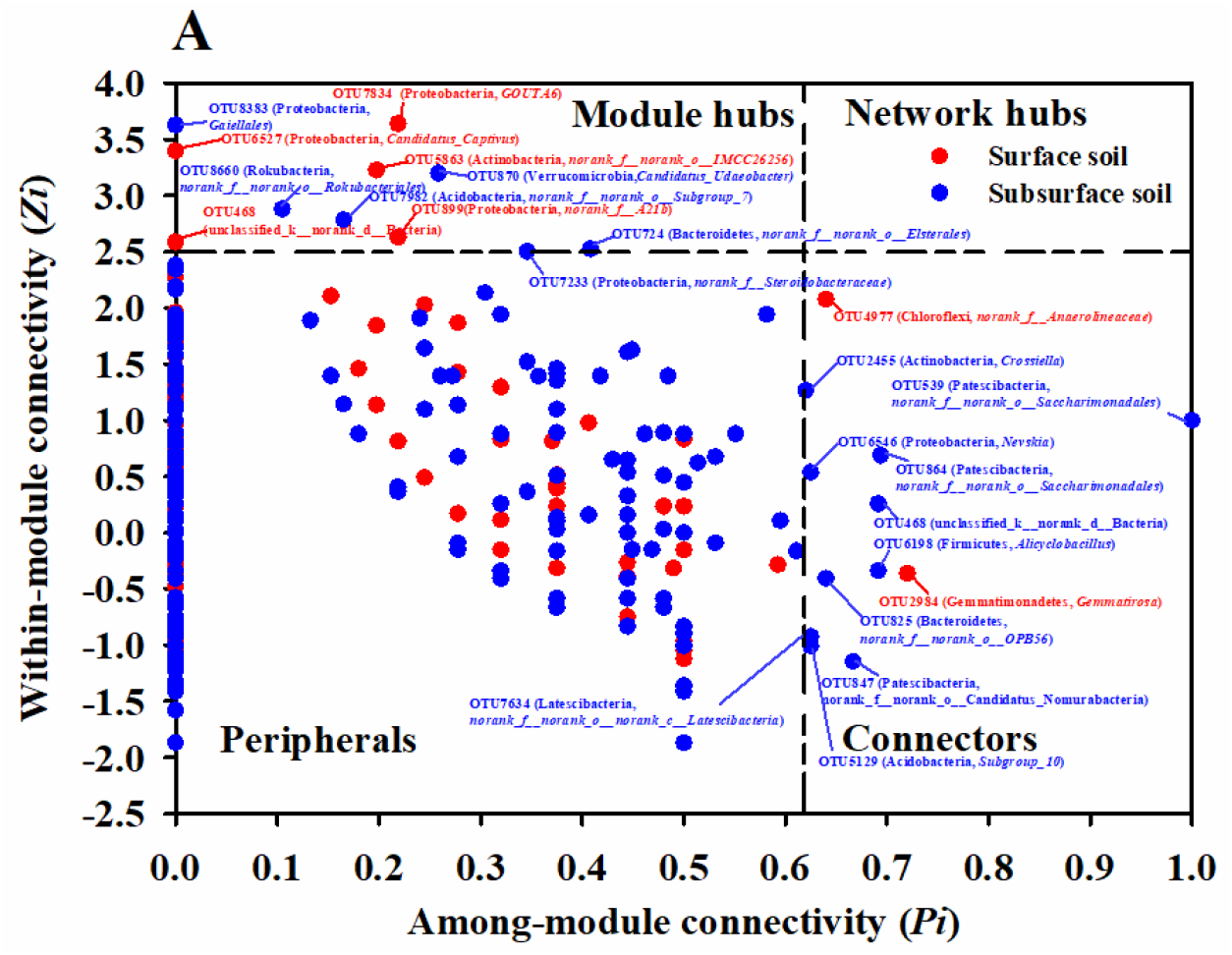
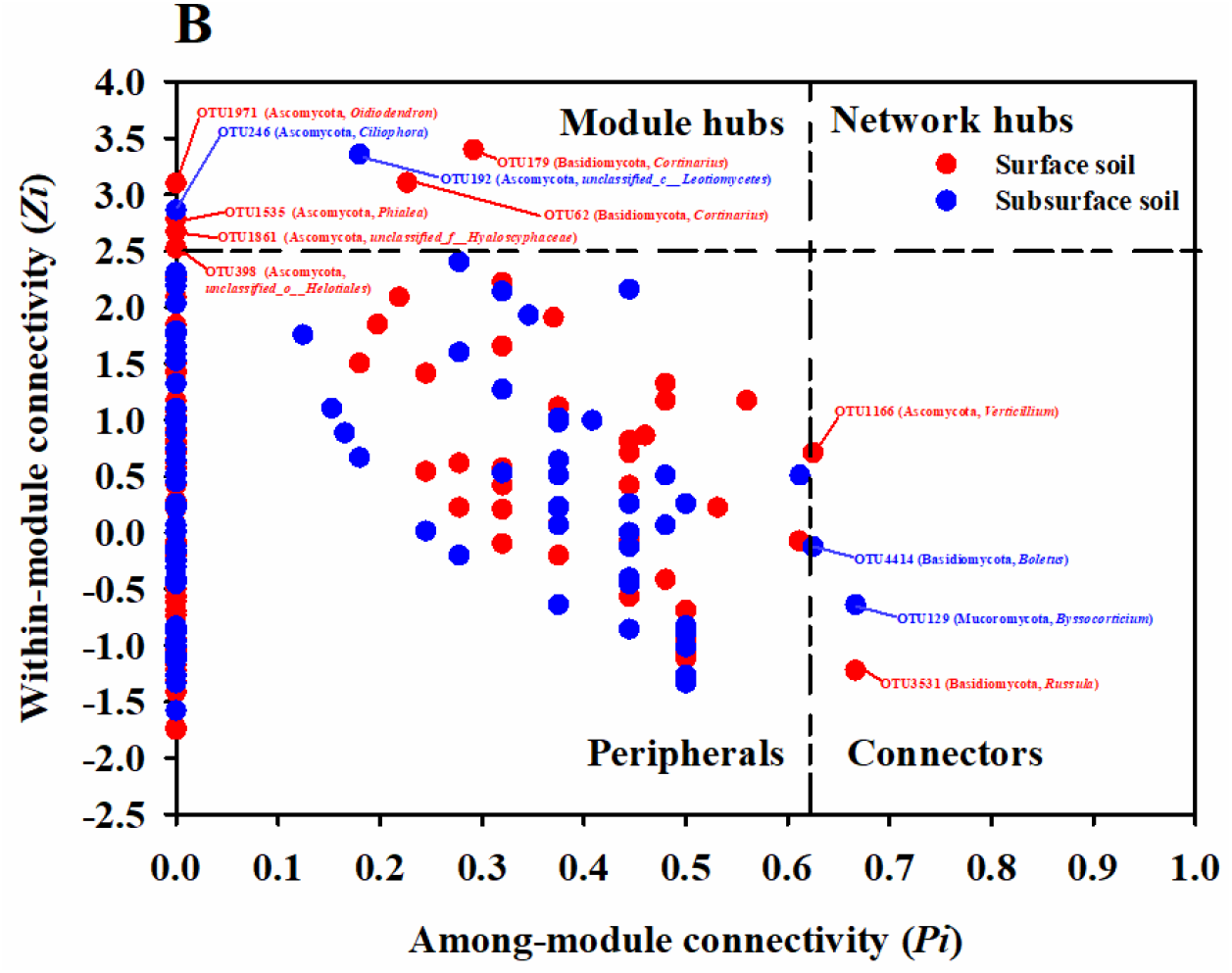
Topological roles of OTUs in the soil bacterial (A) and fungal (B) co-occurrence networks as indicated by the Zi-Pi plot. The nodes with Zi > 2.5 are identified as module hubs, and those with Pi > 0.62 are connectors. The network hubs are determined by Zi > 2.5 and Pi > 0.62, and the peripherals are characterized by Zi < 2.5 and Pi < 0.62

## 4 Discussion

Our results highlighted several key findings related to the altitude distribution of soil bacterial and fungal communities in cold temperate zones. Firstly, similar to the temperate and tropical climate conditions, the bacterial and fungal communities in the cold-temperate mountain ecosystems showed inconsistent patterns, that is, monotonous declining and monotonic increasing, respectively. Then, compared to the soil depth, the bacterial and fungal community structure was more sensitive and fragile to altitude, and the variation of abiotic factors along the altitudinal gradient dominated the changes in the microbial community. Finally, the co-occurrence network of bacteria in the subsurface soil had high complexity and modularity, while the complexity of the fungal network did not change with the increasing soil depth.

### 4.1 Divergent controlling factors for bacterial and fungal diversity and community composition along an altitudinal gradient

Previous studies of microbial diversity in mountain ecosystems reported different altitude-diversity patterns (Shen et al., 2013; Shen et al., 2014; Singh et al., 2014; Peay et al., 2017; Ren et al., 2018; Guo et al., 2020; Shen et al., 2020). Similar to the results of most studies based on high-throughput sequencing technology (Li et al., 2018; Shen et al., 2015; Shen et al., 2019), we found that soil bacterial diversity decreased with increasing altitude, however, the fungal diversity increased with altitude, which partially supported our first hypothesis. Some recent studies also have emphasized the inconsistency of bacterial and fungal biogeographical patterns (Peay et al., 2017; Bahram et al., 2018; Shen et al., 2020). Peay et al. (2017) pointed out that due to the large differences in the life and evolutionary histories of different taxa, soil bacteria (single peak) and fungi (linear increase) in the Mt. Hawaiian show different altitude distribution patterns. In general, the harsh degree of the environment increases with altitude, so that it is expected that the abundance of bacteria and fungi will decrease along the altitudinal gradient (Margesin et al., 2009). However, we found that soil fungi maintained a higher diversity at high altitudinal samples, which may be due to higher soil nutrient levels (DON and ammonium nitrogen), which in turn promoted the growth of microorganisms (Peay et al., 2017). In addition, we found that the diversity of bacterial communities was higher than that of fungal communities, which is consistent with the study of Meng et al. (2013) in subtropical mountain ecosystems, this result implied the niche differentiation of different microbial groups along the altitudinal gradient in the cold temperate zone (Prosser et al., 2007). In this study, the microbial abundance of different taxa showed different responses to altitude and soil layer. Altitude had a marked effect on some of the higher abundance bacterial phyla (Actinobacteria, Chloroflexi, Planctomycetes) and fungal classes (Agaricomycetes, Leotiomycetes, Pezizomycetes, Umbelopsidomycetes). Shen et al. (2020) recently conducted a more fine-resolution comparison of the diversity of bacterial and fungal communities in the Mt. Kilimanjaro in East Africa, and pointed out that the diversity patterns of taxonomic groups (phyla or classes) in bacterial and fungal communities were different and same, respectively. Due to the uneven distribution of microbial effective nutrients and plant roots along the soil profile, the contribution of soil depth may be higher than the geographic difference of soil microbial communities (Rousk et al., 2010). In this study, whether it was soil bacteria or fungi, there was no significant difference in their community diversity between the surface layer and the subsurface layer. The soil depth had a statistically significant impact on the bacterial community richness, but had no significant effect on fungi. This is probably due to soil fungi have a narrower physiological range than bacteria. For example, soil fungi are heterotrophic organisms, while soil bacteria can be photosynthetic autotrophic organisms, heterotrophic organisms or chemoautotrophic organisms (Lladó et al., 2017). Based on the results of PERMANOVA, we further verified that in the cold-temperate mountain ecosystem, the influence of altitude on the community structure of bacteria and fungi is stronger than that of the soil depth.

As we expected, soil pH in cold-temperate mountain ecosystems was a good predictor of soil bacterial community composition, which is consistent with most earlier studies (Shen et al., 2013; Bahram et al., 2018; Shen et al., 2019), the pH range in this study (3.92~4.74) was similar to the study in Changbai Mountain (3.89~6.31) (Shen et al., 2013), although our pH variability was very small. A previous study reported the effect of varying soil pH in a small range on bacterial community structure (Sagova-Mareckova et al., 2015). Rousk et al. (2010) pointed out that the composition of bacterial communities was mainly affected by soil pH, not due to diffusion limitations between microbial communities and other environmental factors. Although many studies had reported the relationship between pH and bacterial community composition and diversity, in our study, except soil pH, soil moisture also played an important role in bacterial community. The study of Shen et al. (2020) in Mt. Kilimanjaro pointed out that the average annual rainfall was the second most important factor in predicting soil bacterial diversity, which indirectly affected bacterial communities by regulating pH and plant productivity (Tian et al., 2018). Although many studies had reported the relationship between temperature and soil fungal communities (Jarvis et al., 2015; Newsham et al., 2016; Shen et al., 2020), our results weren’t line with our expectation that temperature was the main factor affecting the diversity and composition of the fungal community. In our study, DON played a critical role in affecting the composition of soil fungal community, followed by NH_4_^+^-N. To our knowledge, this was the first reported observation that DON was an important factor in predicting the variation of soil fungal community composition and diversity along an altitudinal gradient. Dissolved organic matter (DOM) is an important part of soil organic matter and provides organic substrates and resources for heterotrophic microorganisms (Benner, 2011; Huang et al., 2020). Huang et al. (2020) found in a recent study that DOM quality was the most important driving factor explaining the diversity and community composition of soil fungi. In this study, altitude has a significant impact on DON. Vegetation types among different altitudes have specific effects on the soil physical and chemical properties, especially through litter pathways that lead to certain differences in the composition of soil organic matter (Quideau et al., 2001), resulting in different soil microclimates (Knelman et al., 2012). Shen et al. (2016) reported that DOC can predict the functional genetic diversity of microorganisms in the Changbai Mountain ecosystem from forest to tundra. Also, in the study of the small-scale altitudinal gradient of the Changbai Mountain tundra, Ni et al. (2018) found that the abundance of Ascomycota and ectomycorrhizal fungi were significantly correlated with the content of DON and NH_4_^+^-N, respectively. In general, our research highlighted the different driving factors for the altitude distribution of bacterial and fungal communities in cold-temperate forest soils.

### 4.2 Potentially more connected network of soil bacteria in surface soil than that in subsurface soils

The microbial community is a complex combination of highly interactive taxa (Fuhrman, 2009). Understanding the correlation of microbes is essential for predicting the response of microbial communities to climate change, microbial co-occurrence networks with lower complexity are easily considered stressed by the environment (Banerjee et al., 2019). It is worth noted that despite the soil depth had a small effect on the composition and diversity of bacterial and fungal communities, the co-occurrence network of bacteria and fungi showed different response patterns to the soil depth, which was consistent with our third hypothesis. For bacterial community, the differences of the network between different soil layers were more obvious, that is, the network of subsurface soil had greater modularity, density and more highly connected nodes than the surface layer. In contrast, there was no obvious difference in the fungal co-occurrence network between different soil layers. To the best of our knowledge, this is the first reported study of the co-occurrence network of microorganisms along the soil profile in the cold temperate zone of China. In a recent study, de Vries et al. (2018) found that the network of soil fungi was more stable in response to extreme conditions than bacteria, in addition to vegetation composition, soil moisture played a key role. In this study, the soil moisture and temperature were highly variable along the soil profile, which may lead to the difference in the response of this co-occurring network of different microbial groups to the two contrasting soil layers. A recent study by Tu et al. (2020) on six forests in the United States found that soil temperature and soil water content were highly correlated with the modularity of the microbial co-occurrence network. In addition, a possible mechanism behind the more connected network was the reduction of root input, metabolites and the number of available substrates in the subsurface soil, which caused more competition or co-metabolism for substrates of a wide variety of bacterial communities (Upton et al., 2020). Compared with the fungal network topology, the bacterial network was more complex, which also implied that the bacterial communities in the cold-temperate mountain ecosystem were more sensitive to the variation of environmental factors along the soil profiles. Different from our results, Xiao et al. (2018) compared the *Phyllostachys edulis* plantation and pointed out that the degree of connectivity of the bacterial network was lower than that of fungi, which might imply that the different interaction pattern of microorganisms varied between different habitats. In this study, OTUs belonging to Proteobacteria and Actinobacteria were mainly used as modular hubs and network connectors of bacterial networks, playing a critical role in bacterial co-occurrence networks between different soil layers. Proteobacteria is usually the dominant nitrogen-fixing bacteria phylum in soil ecosystems (Gaby and Buckley, 2011). Actinobacteria has a mycelial growth pattern in the soil, allowing plants to expand the surface area in a deeper soil layer to absorb nutrients, and may become the soil aggregate and potentially active components that preserve water and nutrients (Fierer et al., 2013; Upton et al., 2020). However, the bacterial networks in the surface and subsurface layers had different keystone compositions, which further confirmed the niche differentiation of bacterial taxonomic species along the soil profiles. Our results implied the different patterns of bacterial and fungal networks along the soil profiles in cold-temperate mountain ecosystems.

## 5 Conclusion

Our research described for the first time the biogeographic distribution of soil microbial communities in the cold-temperate mountain ecosystem in China. Our results confirmed that soil bacterial (monotonously decreasing) and fungal (monotonously increasing) diversity showed inconsistent altitude distribution patterns. The dramatic variations in soil factors along the altitudinal gradient were the main causes driving the variation in the community composition and diversity of bacteria (pH) and fungi (DON). Although the soil microbial community was more affected by the altitudinal gradient than the soil depth, the network analysis further emphasized the obvious differences in the bacterial and fungal communities between the two contrasting soil layers. Compared with soil fungi, soil bacterial communities were more sensitive to changes in soil quality along the soil profile, and bacterial networks in subsurface soils exhibit more complex and compact topological features. Further research could focus on specific taxa, microbial interactions, and the functions of keystones in a forest ecosystem. This is essential for a better understanding of the mechanisms that affect microbial diversity and functions in this fragile ecosystem.

## Reference

Bahram, M., Hildebrand, F., Forslund, S.K., Anderson, J.L., Soudzilovskaia, N.A., Bodegom, P.M., Bengtsson-Palme, J., Anslan, S., Coelho, L.P., Harend, H., 2018. Structure and function of the global topsoil microbiome. Nature 560, 233–237.

Bahram, M., Polme, S., Koljalg, U., Zarre, S., Tedersoo, L., 2012. Regional and local patterns of ectomycorrhizal fungal diversity and community structure along an altitudinal gradient in the Hyrcanian forests of northern Iran. New Phytol. 193, 465–473.

Baldrian, P., 2017. Forest microbiome: diversity, complexity and dynamics. FEMS Microbiol. Rev. 41, 109–130.

Banerjee, S., Walder, F., Buchi, L., Meyer, M., Held, A.Y., Gattinger, A., Keller, T., Charles, R., Der Heijden, M.G.A.V., 2019. Agricultural intensification reduces microbial network complexity and the abundance of keystone taxa in roots. ISME J. 13, 1722–1736.

Banning, N.C., Gleeson, D.B., Grigg, A.H., Grant, C.D., Andersen, G.L., Brodie, E.L., Murphy, D., 2011. Soil microbial community successional patterns during forest ecosystem restoration. Appl. Environ. Microbiol. 77, 6158–6164.

Benner, R., 2011. Biosequestration of carbon by heterotrophic microorganisms. Nat. Rev. Microbiol. 9, 75–75.

Brookes, P., Landman, A., Pruden, G., Jenkinson, D., 1985. Chloroform fumigation and the release of soil nitrogen: a rapid direct extraction method to measure microbial biomass nitrogen in soil. Soil Biol. Biochem. 17, 837–842.

Brown, J.H., Gillooly, J.F., Allen, A.P., Savage, V.M., West, G.B., 2004. Toward a metabolic theory of ecology. Ecology 85, 1771–1789.

Bryant, J.A., Lamanna, C., Morlon, H., Kerkhoff, A.J., Enquist, B.J., Green, J.L., 2008. Colloquium paper: microbes on mountainsides: contrasting elevational patterns of bacterial and plant diversity. Proc. Natl. Acad. Sci. Unit. States Am. 105 Suppl 1, 11505–11511.

Caporaso, J.G., Lauber, C.L., Walters, W.A., Berglyons, D., Huntley, J., Fierer, N., Owens, S.M., Betley, J.R., Fraser, L., Bauer, M., 2012. Ultra-high-throughput microbial community analysis on the Illumina HiSeq and MiSeq platforms. ISME J. 6, 1621–1624.

Cardona, C., Weisenhorn, P., Henry, C., Gilbert, J.A., 2016. Network-based metabolic analysis and microbial community modeling. Curr. Opin. Microbiol. 31, 124–131.

Christensen, T.R., Johansson, T., Åkerman, H.J., Mastepanov, M., Malmer, N., Friborg, T., Crill, P., Svensson, B.H., 2004. Thawing sub arctic permafrost: Effects on vegetation and methane emissions. Geophys. Res. Lett. 31.

Cline, M.S., Smoot, M., Cerami, E., Kuchinsky, A., Landys, N., Workman, C., Christmas, R., Avila-Campilo, I., Creech, M., Gross, B., 2007. Integration of biological networks and gene expression data using Cytoscape. Nat. Protoc. 2, 2366.

Coolen, M.J., van de Giessen, J., Zhu, E.Y., Wuchter, C., 2011. Bioavailability of soil organic matter and microbial community dynamics upon permafrost thaw. Environ. Microbiol. 13, 2299–2314.

de Vries, F.T., Griffiths, R.I., Bailey, M., Craig, H., Girlanda, M., Gweon, H.S., Hallin, S., Kaisermann, A., Keith, A.M., Kretzschmar, M., 2018. Soil bacterial networks are less stable under drought than fungal networks. Nat. Commun. 9, 3033.

Deng, J., Gu, Y., Zhang, J., Xue, K., Qin, Y., Yuan, M., Yin, H., He, Z., Wu, L., Schuur, E.A., 2015. Shifts of tundra bacterial and archaeal communities along a permafrost thaw gradient in A laska. Mol. Ecol. 24, 222–234.

Du, X., Deng, Y., Li, S., Escalas, A., Feng, K., He, Q., Wang, Z., Wu, Y., Wang, D., Peng, X., 2021. Steeper spatial scaling patterns of subsoil microbiota are shaped by deterministic assembly process. Mol. Ecol. 30, 1072–1085.

Edgar, R.C., 2013. UPARSE: highly accurate OTU sequences from microbial amplicon reads. Nat. Method. 10, 996–998.

Eilers, K.G., Debenport, S., Anderson, S., Fierer, N., 2012. Digging deeper to find unique microbial communities: the strong effect of depth on the structure of bacterial and archaeal communities in soil. Soil Biol. Biochem. 50, 58–65.

Fan, K., Cardona, C., Li, Y., Shi, Y., Xiang, X., Shen, C., Wang, H., Gilbert, J.A., Chu, H., 2017. Rhizosphere-associated bacterial network structure and spatial distribution differ significantly from bulk soil in wheat crop fields. Soil Biol. Biochem. 113, 275–284.

Faust, K., Raes, J., 2012. Microbial interactions: from networks to models. Nat. Rev. Microbiol. 10, 538–550.

Fierer, N., Ladau, J., Clemente, J.C., Leff, J.W., Owens, S.M., Pollard, K.S., Knight, R., Gilbert, J.A., McCulley, R.L., 2013. Reconstructing the microbial diversity and function of pre-agricultural tallgrass prairie soils in the United States. Science 342, 621–624.

Fierer, N., McCain, C.M., Meir, P., Zimmermann, M., Rapp, J.M., Silman, M.R., Knight, R., 2011. Microbes do not follow the elevational diversity patterns of plants and animals. Ecology 92, 797–804.

Fritze, H., Pietikäinen, J., Pennanen, T., 2000. Distribution of microbial biomass and phospholipid fatty acids in Podzol profiles under coniferous forest. Eur. J. Soil Sci. 51, 565–573.

Fuhrman, J.A., 2009. Microbial community structure and its functional implications. Nature 459, 193–199.

Gaby, J.C., Buckley, D.H., 2011. A global census of nitrogenase diversity. Environ. Microbiol. 13, 1790–1799.

Gade, L., Scheel, C.M., Pham, C.D., Lindsley, M.D., Iqbal, N., Cleveland, A.A., Whitney, A.M., Lockhart, S.R., Brandt, M.E., Litvintseva, A.P., 2013. Detection of fungal DNA in human body fluids and tissues during a multistate outbreak of fungal meningitis and other infections. Eukaryot. Cell 12, 677–683.

Guo, Y., Ren, C., Yi, J., Doughty, R., Zhao, F., 2020. Contrasting responses of rhizosphere bacteria, fungi and arbuscular mycorrhizal fungi along an elevational gradient in a temperate montane forest of China. Front. Microbiol. 11, 2042.

Hannula, S.E., Morrien, E., de Hollander, M., van der Putten, W.H., van Veen, J.A., de Boer, W., 2017. Shifts in rhizosphere fungal community during secondary succession following abandonment from agriculture. ISME J. 11, 2294–2304.

Hawkins, B.A., Field, R., Cornell, H.V., Currie, D.J., Guégan, J.-F., Kaufman, D.M., Kerr, J.T., Mittelbach, G.G., Oberdorff, T., O’Brien, E.M., 2003. Energy, water, and broad? scale geographic patterns of species richness. Ecology 84, 3105–3117.

Huang, M., Chai, L., Jiang, D., Zhang, M., Jia, W., Huang, Y., 2020. Spatial patterns of soil fungal communities are driven by dissolved organic matter (DOM) quality in semi-arid regions. Microb. Ecol. 1–913.

Jarvis, S.G., Woodward, S., Taylor, A.F., 2015. Strong altitudinal partitioning in the distributions of ectomycorrhizal fungi along a short (300 m) elevation gradient. New Phytol. 206, 1145–1155.

Joergensen, R.G., 1996. The fumigation-extraction method to estimate soil microbial biomass: calibration of the kEC value. Soil Biol. Biochem. 28, 25–31.

Kirchman, D.L., 2018. Processes in microbial ecology. Oxford University Press.

Knelman, J.E., Legg, T.M., O’Neill, S.P., Washenberger, C.L., González, A., Cleveland, C.C., Nemergut, D.R., 2012. Bacterial community structure and function change in association with colonizer plants during early primary succession in a glacier forefield. Soil Biol. Biochem. 46, 172–180.

Lee, C.K., Barbier, B.A., Bottos, E.M., McDonald, I.R., Cary, S.C., 2012. The inter-valley soil comparative survey: the ecology of Dry Valley edaphic microbial communities. ISME J. 6, 1046–1057.

Li, G., Xu, G., Shen, C., Tang, Y., Zhang, Y., Ma, K., 2016. Contrasting elevational diversity patterns for soil bacteria between two ecosystems divided by the treeline. Sci. China Life Sci. 59, 1177–1186.

Li, J., Li, C., Kou, Y., Yao, M., He, Z., Li, X., 2020. Distinct mechanisms shape soil bacterial and fungal co-occurrence networks in a mountain ecosystem. FEMS Microbiol. Ecol. 96.

Li, J., Shen, Z., Li, C., Kou, Y., Wang, Y., Tu, B., Zhang, S., Li, X., 2018. Stair-step pattern of soil bacterial diversity mainly driven by ph and vegetation types along the elevational gradients of gongga mountain, China. Front. Microbiol. 9, 569.

Lladó, S., López-Mondéjar, R., Baldrian, P., 2017. Forest soil bacteria: diversity, involvement in ecosystem processes, and response to global change. Microbiol. Mol. Biol. Rev. 81.

Luan, L., Liang, C., Chen, L., Wang, H., Xu, Q., Jiang, Y., Sun, B., 2020. Coupling Bacterial Community Assembly to Microbial Metabolism across Soil Profiles. mSystems 5.

Ma, B., Dai, Z., Wang, H., Dsouza, M., Liu, X., He, Y., Wu, J., Rodrigues, J.L., Gilbert, J.A., Brookes, P.C., 2017. Distinct biogeographic patterns for archaea, bacteria, and fungi along the vegetation gradient at the continental scale in Eastern China. Msystems 2.

Margesin, R., Jud, M., Tscherko, D., Schinner, F., 2009. Microbial communities and activities in alpine and subalpine soils. FEMS Microbiol. Ecol. 67, 208–218.

Meng, H., Li, K., Nie, M., Wan, J.-R., Quan, Z.-X., Fang, C.-M., Chen, J.-K., Gu, J.-D., Li, B., 2013. Responses of bacterial and fungal communities to an elevation gradient in a subtropical montane forest of China. Appl. Microbiol. Biotechnol. 97, 2219–2230.

Miyamoto, Y., Nakano, T., Hattori, M., Nara, K., 2014. The mid-domain effect in ectomycorrhizal fungi: range overlap along an elevation gradient on Mount Fuji, Japan. ISME J. 8, 1739–1746.

Newsham, K.K., Hopkins, D.W., Carvalhais, L.C., Fretwell, P.T., Rushton, S.P., O’Donnell, A.G., Dennis, P.G., 2016. Relationship between soil fungal diversity and temperature in the maritime Antarctic. Nat. Clim. Change 6, 182–186.

Ni, Y., Yang, T., Zhang, K., Shen, C., Chu, H., 2018. Fungal communities along a small-scale elevational gradient in an alpine tundra are determined by soil carbon nitrogen ratios. Front. Microbiol. 9, 1815.

Peay, K.G., von Sperber, C., Cardarelli, E., Toju, H., Francis, C.A., Chadwick, O.A., Vitousek, P.M., 2017. Convergence and contrast in the community structure of bacteria, fungi and archaea along a tropical elevation–climate gradient. FEMS Microbiol. Ecol. 93.

Prosser, J.I., Bohannan, B.J., Curtis, T.P., Ellis, R.J., Firestone, M.K., Freckleton, R.P., Green, J.L., Green, L.E., Killham, K., Lennon, J.J., 2007. The role of ecological theory in microbial ecology. Nat. Rev. Microbiol. 5, 384–392.

Quideau, S., Chadwick, O., Benesi, A., Graham, R., Anderson, M., 2001. A direct link between forest vegetation type and soil organic matter composition. Geoderma 104, 41–60.

Reich, P.B., 2012. Key canopy traits drive forest productivity. Proc. Royal Soc. B: Biol.Sci. 279, 2128–2134.

Reich, P.B., Frelich, L.E., Voldseth, R.A., Bakken, P., Adair, E.C., 2012. Understorey diversity in southern boreal forests is regulated by productivity and its indirect impacts on resource availability and heterogeneity. J. Ecol. 100, 539–545.

Ren, C., Zhang, W., Zhong, Z., Han, X., Yang, G., Feng, Y., Ren, G., 2018. Differential responses of soil microbial biomass, diversity, and compositions to altitudinal gradients depend on plant and soil characteristics. Sci. Total Environ. 610-611, 750.

Rousk, J., Bååth, E., 2007. Fungal biomass production and turnover in soil estimated using the acetate-in-ergosterol technique. Soil Biol. Biochem. 39, 2173–2177.

Rousk, J., Bååth, E., Brookes, P.C., Lauber, C.L., Lozupone, C., Caporaso, J.G., Knight, R., Fierer, N., 2010. Soil bacterial and fungal communities across a pH gradient in an arable soil. ISME J. 4, 1340–1351.

Sagova-Mareckova, M., Cermak, L., Omelka, M., Kyselkova, M., Kopecky, J., 2015. Bacterial diversity and abundance of a creek valley sites reflected soil pH and season. Open Life Sci. 1.

Seidl, R., Albrich, K., Erb, K., Formayer, H., Leidinger, D., Leitinger, G., Tappeiner, U., Tasser, E., Rammer, W., 2019. What drives the future supply of regulating ecosystem services in a mountain forest landscape? Forest Ecol. Manag. 445, 37–47.

Shen, C., Gunina, A., Luo, Y., Wang, J., He, J.Z., Kuzyakov, Y., Hemp, A., Classen, A.T., Ge, Y., 2020. Contrasting patterns and drivers of soil bacterial and fungal diversity across a mountain gradient. Environ. Microbiol.

Shen, C., Liang, W., Shi, Y., Lin, X., Zhang, H., Wu, X., Xie, G., Chain, P., Grogan, P., Chu, H., 2014. Contrasting elevational diversity patterns between eukaryotic soil microbes and plants. Ecology 95, 3190–3202.

Shen, C., Ni, Y., Liang, W., Wang, J., Chu, H., 2015. Distinct soil bacterial communities along a small-scale elevational gradient in alpine tundra. Front. Microbiol. 6, 582.

Shen, C., Shi, Y., Fan, K., He, J., Adams, J.M., Ge, Y., Chu, H., 2019. Soil pH dominates elevational diversity pattern for bacteria in high elevation alkaline soils on the Tibetan Plateau. FEMS Microbiol. Ecol. 95.

Shen, C., Shi, Y., Ni, Y., Deng, Y., Van Nostrand, J.D., He, Z., Zhou, J., Chu, H., 2016. Dramatic increases of soil microbial functional gene diversity at the treeline ecotone of Changbai mountain. Front. Microbiol. 7, 1184.

Shen, C., Xiong, J., Zhang, H., Feng, Y., Lin, X., Li, X., Liang, W., Chu, H., 2013. Soil pH drives the spatial distribution of bacterial communities along elevation on Changbai Mountain. Soil Biol. Biochem. 57, 204–211.

Sheng, Y., Cong, J., Lu, H., Yang, L., Liu, Q., Li, D., Zhang, Y., 2019. Broad leaved forest types affect soil fungal community structure and soil organic carbon contents. MicrobiologyOpen 8, e874.

Singh, D., Lee-Cruz, L., Kim, W.-S., Kerfahi, D., Chun, J.-H., Adams, J.M., 2014. Strong elevational trends in soil bacterial community composition on Mt. Halla, South Korea. Soil Biol. Biochem. 68, 140–149.

Sundqvist, M.K., Sanders, N.J., Wardle, D.A., 2013. Community and ecosystem responses to elevational gradients: processes, mechanisms, and insights for global change. Annu. Rev. Ecol. Evol. Syst. 44, 261–280.

Team, R.C., 2013. R: A language and environment for statistical computing.

Tian, J., He, N., Hale, L., Niu, S., Yu, G., Liu, Y., Blagodatskaya, E., Kuzyakov, Y., Gao, Q., Zhou, J., 2018. Soil organic matter availability and climate drive latitudinal patterns in bacterial diversity from tropical to cold temperate forests. Funct. Ecol. 32, 61–70.

Tu, Q., Yan, Q., Deng, Y., Michaletz, S.T., Buzzard, V., Weiser, M.D., Waide, R., Ning, D., Wu, L., He, Z., 2020. Biogeographic patterns of microbial co-occurrence ecological networks in six American forests. Soil Biol.Biochem. 107897.

Upton, R.N., Checinska Sielaff, A., Hofmockel, K.S., Xu, X., Polley, H.W., Wilsey, B.J., 2020. Soil depth and grassland origin cooperatively shape microbial community co occurrence and function. Ecosphere 11, e02973.

Widder, S., Allen, R.J., Pfeiffer, T., Curtis, T.P., Wiuf, C., Sloan, W.T., Cordero, O.X., Brown, S.P., Momeni, B., Shou, W., 2016. Challenges in microbial ecology: building predictive understanding of community function and dynamics. ISME J. 10, 2557–2568.

Xiao, X., Liang, Y., Zhou, S., Zhuang, S., Sun, B., 2018. Fungal community reveals less dispersal limitation and potentially more connected network than that of bacteria in bamboo forest soils. Mol. Ecol. 27, 550–563.

Yang, T., Adams, J.M., Shi, Y., Sun, H., Cheng, L., Zhang, Y., Chu, H., 2017. Fungal community assemblages in a high elevation desert environment: Absence of dispersal limitation and edaphic effects in surface soil. Soil Biol. Biochem. 115, 393–402.

Zhou, J., Deng, Y., Shen, L., Wen, C., Yan, Q., Ning, D., Qin, Y., Xue, K., Wu, L., He, Z., 2016. Temperature mediates continental-scale diversity of microbes in forest soils. Nat. Commun. 7, 1–10.

