## Supplemental files for "Contrasting patterns and co-occurrence network of soil bacterial and fungal community along depth profiles in cold temperate montane forests of China"

Lixue Yang;

**Running Title:** Contrasting altitudinal patterns of soil bacteria and fungi

**Article Type:** Original research paper

**Number of words:** 6277

**Number of text pages:** 24

**Number of tables and figures:** 3 tables and 7 figures

**Number of references:** 74

**Supplementary material captions**

**Supplementary Table 1** Site characteristics of different altitudes

**Supplementary Table 2** Two-way analysis of variance of relative abundance of dominant bacterial phyla and classes

**Supplementary Table 3** Two-way analysis of variance of relative abundance of dominant fungal phyla and classes

**Supplementary Table 4** Non-parametric multivariate analysis (PERMANOVA) of soil bacterial and fungal community by altitude and soil depth

**Supplementary Figure 1** Location of the Oakley Mountains in Greater Khingan Mountains

**Supplementary Figure 2** The Rarefaction curves of the number of operational taxonomic units (OTUs) for soil bacterial (A) and fungal (B) communities.

**Supplementary Table 1 Site characteristics of different altitudes**

| Altitude  (m, a.s.l.) | Coordinates | Soil type | Vegetation type | Dominant taxa |
| --- | --- | --- | --- | --- |
| 830 | N 51°47′41″  E 122°5′3″ | Dark brown coniferous soil | Cold temperate coniferous forest | *Larix gmelinii*、*Vaccinium vitis-idaea*、*Ledum palustre*、*Rosa davurica*、*Lonicera caerulea*、*Rubus arcticus*、*Pyrola incarnata*、*Deyeuxia angustifolia* |
| 950 | N 51°49′42″  E 122°3′34″ | Dark brown coniferous soil | Cold temperate coniferous forest | *Larix gmelinii*、*Betula platyphylla*、*Vaccinium vitis-idaea*、*Ledum palustre*、*Rhododendron dauricum*、*Spiraea dahurica*、*Rubus sachalinensis*、*Sambucus williamsii*、*Rosa davurica*、*Artemisia lagocephala*、*Cimicifuga foetida*、*Vicia ramuliflora*、*Pyrola incarnata* |
| 1100 | N 51°49′89″  E 122°2′76″ | Podzolic brown coniferous soil | Cold temperate coniferous forest | *Larix gmelinii*、*Pinus sylvestris*、*Pinus pumila*、*Betula platyphylla*、*Vaccinium vitis-idaea*、*Ledum palustre*、*Rhododendron dauricum*、*Spiraea dahurica*、*Rubus sachalinensis*、*Sambucus williamsii*、*Artemisia lagocephala*、*Clematis sibirica*、*Cimicifuga foetida*、*Vicia ramuliflora* |
| 1300 | N 51°50′14″  E 122°2′19″ | Podzolic brown coniferous soil | Cold temperate coniferous forest | *Larix gmelinii*、*Pinus pumila*、*Betula ermanii*、*Rhododendron dauricum*、*Vaccinium vitis-idaea*、*Ledum palustre*、*Artemisia lagocephala*、*Aquilegia viridiflora*、*Saxifraga bronchialis*、*Polygonum alpinum* |

a.s.l. = above sea level

**Supplementary Table 2 Two-way analysis of variance of relative abundance of dominant bacterial phyla and classes**

| Taxonomy | | Altitude | | Soil depth | | Altitude×Soil depth | |
| --- | --- | --- | --- | --- | --- | --- | --- |
|  |  | F | *P* | F | *P* | F | *P* |
| Phyla | Proteobacteria | 3.146 | 0.054 | 0.213 | 0.651 | 1.236 | 0.329 |
|  | Acidobacteria | 1.431 | 0.271 | 1.276 | 0.275 | 0.084 | 0.968 |
|  | Actinobacteria | 4.928 | **0.013** | 0.965 | 0.341 | 0.390 | 0.762 |
|  | Chloroflexi | 61.919 | **<0.001** | 19.336 | **<0.001** | 0.610 | 0.618 |
|  | Planctomycetes | 8.590 | **0.001** | 16.985 | **0.001** | 1.956 | 0.161 |
|  | Verrucomicrobia | 2.161 | 0.133 | 1.003 | 0.331 | 1.961 | 0.161 |
|  | WPS-2 | 2.957 | 0.064 | 0.038 | 0.849 | 0.601 | 0.623 |
|  | Gemmatimonadetes | 3.094 | 0.057 | 26.875 | **<0.001** | 0.453 | 0.719 |
|  | Bacteroidetes | 1.334 | 0.298 | 0.187 | 0.671 | 0.170 | 0.915 |
|  | Patescibacteria | 5.807 | **0.007** | 3.676 | 0.073 | 0.559 | 0.650 |
|  | Firmicutes | 16.033 | **<0.001** | 6.011 | **0.026** | 0.886 | 0.469 |
| Classes | Alphaproteobacteria | 1.822 | 0.184 | 0.021 | 0.886 | 1.312 | 0.305 |
|  | Acidobacteriia | 3.282 | **0.048** | 2.909 | 0.107 | 0.149 | 0.929 |
|  | Actinobacteria | 4.928 | **0.013** | 0.965 | 0.341 | 0.390 | 0.762 |
|  | AD3 | 29.807 | **<0.001** | 16.862 | **0.001** | 0.240 | 0.867 |
|  | Gammaproteobacteria | 1.946 | 0.163 | 1.403 | 0.253 | 0.060 | 0.980 |
|  | Planctomycetacia | 8.801 | **0.001** | 17.336 | **0.001** | 1.958 | 0.161 |
|  | Verrucomicrobiae | 2.161 | 0.133 | 1.003 | 0.331 | 1.961 | 0.161 |
|  | Deltaproteobacteria | 2.134 | 0.136 | 0.066 | 0.801 | 0.208 | 0.889 |
|  | Subgroup_6 | 0.483 | 0.699 | 0.004 | 0.951 | 1.412 | 0.276 |
|  | norank_p_WPS-2 | 2.957 | 0.064 | 0.038 | 0.849 | 0.601 | 0.623 |
|  | Gemmatimonadetes | 3.094 | 0.057 | 26.875 | **<0.001** | 0.453 | 0.719 |
|  | Bacteroidia | 1.112 | 0.373 | 0.185 | 0.673 | 0.162 | 0.920 |
|  | Ktedonobacteria | 11.820 | **<0.001** | 0.071 | 0.793 | 0.301 | 0.824 |
|  | Saccharimonadia | 3.782 | **0.032** | 6.279 | **0.023** | 0.621 | 0.611 |
|  | TK10 | 61.669 | **<0.001** | 28.633 | **<0.001** | 1.667 | 0.214 |
|  | KD4-96 | 25.943 | **<0.001** | 3.668 | 0.074 | 1.886 | 0.173 |
|  | Bacilli | 19.137 | **<0.001** | 6.165 | **0.024** | 1.004 | 0.417 |
|  | Holophagae | 25.750 | **<0.001** | 31.642 | **<0.001** | 2.966 | 0.063 |
|  | Anaerolineae | 57.998 | **<0.001** | 4.146 | 0.059 | 0.914 | 0.456 |
|  | Blastocatellia_Subgroup_4 | 12.031 | **<0.001** | 4.791 | **0.044** | 2.389 | 0.107 |

**Supplementary Table 3 Two-way analysis of variance of relative abundance of dominant fungal phyla and classes**

| Taxonomy | | Altitude | | Soil depth | | Altitude×Soil depth | |
| --- | --- | --- | --- | --- | --- | --- | --- |
|  |  | F | P | F | P | F | P |
| Phyla | Ascomycota | 4.127 | **0.024** | 0.388 | 0.542 | 1.299 | 0.309 |
|  | Basidiomycota | 4.249 | **0.022** | 0.265 | 0.614 | 2.028 | 0.150 |
|  | Mucoromycota | 3.284 | **0.048** | 2.357 | 0.144 | 1.162 | 0.355 |
|  | Mortierellomycota | 3.232 | 0.050 | 0.090 | 0.767 | 0.241 | 0.867 |
|  | Rozellomycota | 2.201 | 0.128 | 2.570 | 0.128 | 0.788 | 0.518 |
| Classes | Agaricomycetes | 4.838 | **0.014** | 0.107 | 0.747 | 2.473 | 0.099 |
|  | Eurotiomycetes | 0.260 | 0.853 | 0.583 | 0.456 | 0.501 | 0.687 |
|  | Leotiomycetes | 9.501 | **0.001** | 0.250 | 0.624 | 0.608 | 0.619 |
|  | unclassified_p_Ascomycota | 2.448 | 0.101 | 0.195 | 0.664 | 0.898 | 0.464 |
|  | Pezizomycetes | 12.362 | **<0.001** | 0.190 | 0.669 | 0.034 | 0.991 |
|  | Umbelopsidomycetes | 3.394 | **0.044** | 2.301 | 0.149 | 1.212 | 0.338 |
|  | Dothideomycetes | 3.807 | **0.031** | 0.055 | 0.817 | 1.044 | 0.400 |
|  | Archaeorhizomycetes | 0.959 | 0.436 | 0.245 | 0.628 | 0.047 | 0.986 |
|  | Xylonomycetes | 5.949 | **0.006** | 1.722 | 0.208 | 0.597 | 0.626 |
|  | Pezizomycotina_cls_Incertae_sedis | 9.129 | **0.001** | 3.539 | 0.078 | 3.430 | 0.052 |
|  | Mortierellomycetes | 3.234 | 0.050 | 0.091 | 0.767 | 0.239 | 0.868 |
|  | Tritirachiomycetes | 3.968 | **0.027** | 3.563 | 0.077 | 1.279 | 0.315 |
|  | Sordariomycetes | 2.881 | 0.068 | 3.403 | 0.084 | 1.229 | 0.332 |
|  | Saccharomycetes | 3.150 | 0.054 | 2.026 | 0.174 | 0.597 | 0.626 |
|  | Tremellomycetes | 9.458 | **0.001** | 1.049 | 0.321 | 2.981 | 0.063 |
|  | unclassified_p_Rozellomycota | 2.535 | 0.093 | 2.641 | 0.124 | 0.775 | 0.525 |

**Supplementary Table 4 Non-parametric multivariate analysis (PERMANOVA) of soil bacterial and fungal community by altitude and soil depth**

|  |  | Df | Sums of Sqs | Mean Sqs | F.Model | *R^2^* | Pr(>F) |
| --- | --- | --- | --- | --- | --- | --- | --- |
| Bacteria | Altitude | 3 | 0.760 | 0.253 | 7.732 | 0.537 | **0.001** |
|  | Soil depth | 1 | 0.121 | 0.121 | 2.073 | 0.086 | 0.076 |
| Fungi | Altitude | 3 | 2.457 | 0.819 | 7.650 | 0.534 | **0.001** |
|  | Soil depth | 1 | 0.137 | 0.137 | 0.675 | 0.029 | 0.727 |

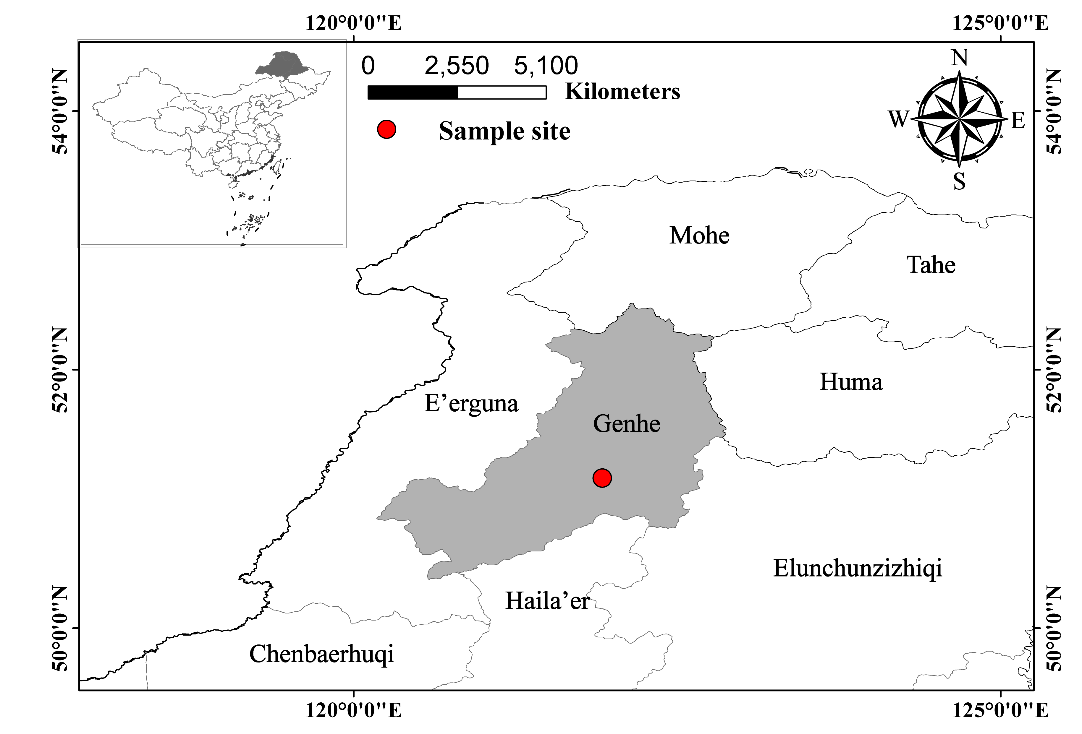

**Supplementary Figure 1 Location of the Oakley Mountains in Greater Khingan Mountains**

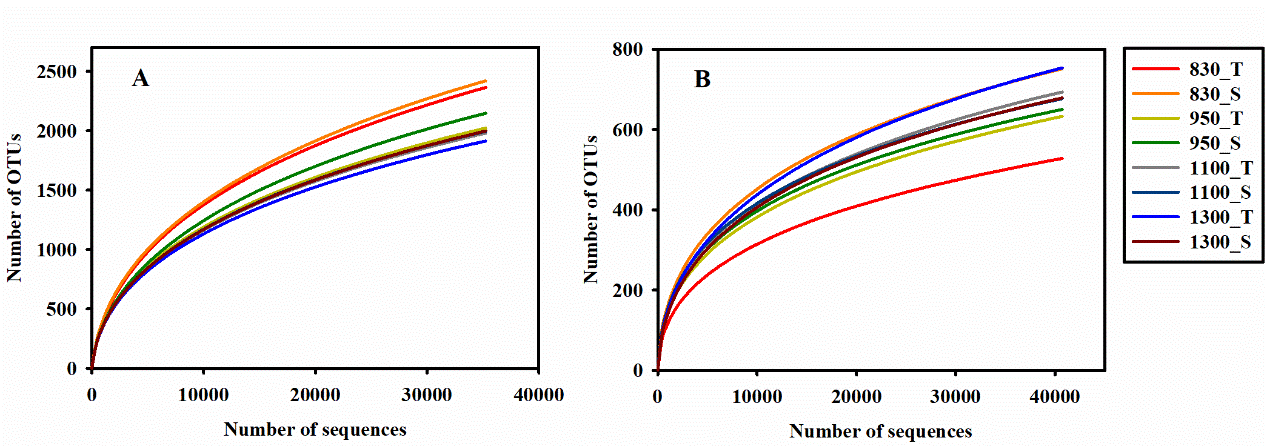

**Supplementary Figure 2 The Rarefaction curves of the number of operational taxonomic units (OTUs) for soil bacterial (A) and fungal (B) communities.** Random subsamples of 49674 and 46434 gene per sample were used to generate the rarefaction curves. OTUs were delineated at 97% sequence similarity. 830_T, 950_T, 1100_T and 1300_T indicate the surface soil in 830 m, 950 m, 1100 m and 1300 m, respectively. 830_S, 950_S, 1100_S and 1300_S indicate the subsurface soil in 830 m, 950 m, 1100 m and 1300 m, respectively.
